# Temporally resolved single-cell RNA sequencing reveals protective and pathological responses during herpes simplex virus 1 CNS infection

**DOI:** 10.1101/2024.08.19.608535

**Authors:** Xiangning Ding, Xin Lai, Ida Hyllen Klæstrup, Sara R. N. Jensen, Morten M Nielsen, Kasper Thorsen, Marina Romero-Ramos, Yonglun Luo, Lin Lin, Line S Reinert, Søren R Paludan

**Affiliations:** Department of Biomedicine, Aarhus University, Aarhus, Denmark; Center for Immunology of Viral Infections, Aarhus University, Aarhus, Denmark; DANDRITE, Aarhus University, Aarhus, Denmark; Department of Molecular Medicine, Aarhus University Hospital, Aarhus, Denmark; Steno Diabetes Center Aarhus, Aarhus University Hospital, Aarhus, Denmark

## Abstract

Herpes Simplex Virus 1 (HSV-1) is a common human neurotropic virus with the majority of adults harboring latent-recurrent infections. In rare cases, HSV-1 infection can access the central nervous system through the neuronal route and develop into life-threatening encephalitis. Here, we used a mouse model for HSV-1 infection to describe the transcriptomic profile of the infected brain stem at the single-cell level and with temporal resolution. Among resident brain cells, microglia increased in proportion during the course of infection, while astrocytes, pericytes, and endothelial cell levels decrease. At the levels of peripheral immune cells, we found notably monocytes to strongly influx the infected brain. Large dynamic changes were found in the abundance of subpopulations of the different cell types following virus infection. For instance, we identify one subpopulation of microglia exhibiting very high type I interferon and chemokine expression early during infection. This population was also enriched for viral transcripts, suggesting localization at foci of infection, and orchestrating recruitment of other immune cells. In contrast, for the infiltrating monocytes, we identified a larger panel of unique subpopulations with antiviral and inflammatory phenotypes, and found not all of these being highly positive for viral transcripts, thus indicating monocyte activities beyond the infected brain areas. Finally, investigation of endothelial cell cross-talk with other cell types revealed that cytokines derived from microglia and monocyte, but also T cells, contribute to disturbance of the blood brain barrier. Our work thus reveals for the first time the complex nature of the cellular response in the virus-infected brain, which seeks to eliminate infection but can also prime for pathological changes.

## Introduction

Herpes simplex encephalitis (HSE) is a severe form of viral encephalitis caused by Herpes Simplex Virus Type 1 (HSV-1) ^1,2^. The disease is characterized by inflammation of the brain tissue, leading to a spectrum of symptoms ranging from mild headaches and fever to severe neurological impairments, such as seizures, memory loss, and potentially life-threatening complications ^1^. HSV-1 is a neurotropic virus that initiate infections in epithelial cells at mucosal surfaces followed by viral infection of sensory neurons in the periphery ^3^. In the majority of cases, this leads to establishment of latent HSV-1 infection in the sensory ganglia. From the state of latency, HSV-1 can periodically be reactivated. However, in rare cases, either during primary infection or reactivation, the infection is not limited to the peripheral nerves, and HSV-1 spreads to the CNS, thus causing encephalitis. Despite advancements in antiviral therapies, HSE continues to pose significant morbidity and mortality, highlighting the critical need for a deeper understanding of its pathophysiological mechanisms.

Viral brain infections are characterized by strong expression of type I interferon (IFN) genes and inflammatory cytokines ^4–6^. This is accompanied by influx of immune cells from the periphery with an early recruitment of monocytes/macrophages and natural killer cells, and later T cells ^7,8^. Both brain resident and infiltrating cell types have been described to contribute to the innate immune response to viral CNS infections. For instance, microglia represent an important early source of IFNβ in the HSV-1-infected mouse brain ^4,9,10^, while in the case of a rhabdovirus infection, this was shown to be mainly derived from astrocytes ^11^. Studies from human genetics and mouse models have shown that defects in the type I IFN system leads to susceptibility to HSE ^12–16^, thus highlighting the importance of this pathway in early host defense in the brain. Use of cell-type specific deficiency in IFN signaling has shown that the action of type I IFNs on neurons and astrocytes is essential to prevent development of disease ^17,18^. Regarding the role of recruited immune cells to the virus-infected brain, it is well-described that CD4 and CD8 T cells play an important part in host defense ^19,20^, largely due to type II IFN-driven antiviral activity ^21^. Studies on how recruited monocytes/macrophages contribute to control of brain infection with viruses, including HSV-1 have given conflicting results ^22–25^.

Since the brain is highly sensitive to immune-mediated pathological responses, understanding of these processes in the context of HSE may provide new treatment options. Studies from various brain inflammation models have shown that inflammatory activity in the brain impair brain development and function. Upon West Nile virus (WNV) encephalitis infection in mice, T cells exert a response dependent on IFN-γ, which leads to microglial activation and cognitive dysfunction after the elimination of virus from the brain ^26^. In the same model, impairment of recruitment of Ly6c+ monocytes into the brain prolonged survival ^24^. In the context of HSV-1, Toll-like receptor 2-deficient mice are less susceptible to acute HSE-like disease, correlating with lower expression of the inflammatory cytokine TNFα ^27^. Data from a human brain organoid model have also identified the TNFα pathway to be highly upregulated upon HSV-1 infection, and revealed that treatment with anti-inflammatory compounds prevented damage of neuronal processes and brain-associated epithelium ^28^. The latter could indicate immune-mediated disruption of the blood-brain-barrier (BBB), which has been observed during WNV infection ^29^. Likewise, detection of a range of markers in the cerebrospinal fluid from HSE patients could indicate BBB break-down ^30^. Thus, the inflammatory response in the virus-infected brain likely impacts on both brain structure and function, but the mechanisms and dynamics are not well understood.

Single cell technologies have emerged as powerful tools to perform deep phenotyping of cellular responses and interactions in complex biological processes *in vivo*. In the context of viral infections, this was first observed in the characterization of COVID-19 pathogenesis, and enabled the identification of how recruited immune cells, notably monocytes turn hyper inflammatory if recruited into a microenvironment with retained viral presence ^31^. With respect to acute viral CNS infections, there is now a panel of studies starting to address the complexities in these diseases with omics technologies ^8,32,33^. For instance, a recent study of brain cells from mice infected with Japanese encephalitis virus (JEV) revealed significant augmentation in communication between individual cells and specific ligand-receptor pairs related to tight junctions and chemokines, indicative of alteration of the BBB and recruitment of immune cells^8^. In one study on HSV-1 infection, CD11b+ immune cells were isolated on day 6 post infection from the ventral posterolateral nucleus thalamic regions and sequenced, which led to identification of a microglia subpopulation enriched for interleukin (IL)-1β signaling, apoptosis, and antigen presentation pathways ^32^.

Collectively, there is lack of detailed information on the host response to HSV-1 infection at the single cell level in the brain at and how this governs the outcome of infection. This includes information on how the responses develop dynamically over time during with respect to both immediate antiviral defense, brain inflammatory response, and interaction with homeostatic brain activities. In this work we have used temporally resolved RNA sequencing of single cells from HSV-1 infected mouse brains to address these questions.

## Result

### HSV-1 CNS infection leads to influx of immune cells and disturbance of brain resident and vasculature cells

To explore the dynamics of transcriptome changes in the HSV-1-infected brain, we inoculated C57BL/6 mice with HSV-1 in the cornea, and isolated material on days 4, 6, and 8 for analysis (Fig. 1a). Infectious virus was detectable in the brain stem on day 4 and 6 post infection, and the mice developed signs of disease, which were observable on days 6 and 8. (Fig. 1b, Extended Data Fig. 1a). To investigate the transcriptomic changes in the microenvironments of the infected mouse brain, we employed GeoMx technology. Different regions of interests (ROIs) exhibiting signs of HSV-1 infection or infiltration of CD45-pos cells in the brain stem or cerebellum were captured and sequenced. (Extended Data Fig. 1b).This analysis revealed that alterations in gene expression was more pronounced in the brainstem compared to the cerebellum following HSV-1 infection, predominantly attributed to the presence of CD45 positive cells (Fig. 1c, Extended Data Fig. 1c).

**Fig. 1.**
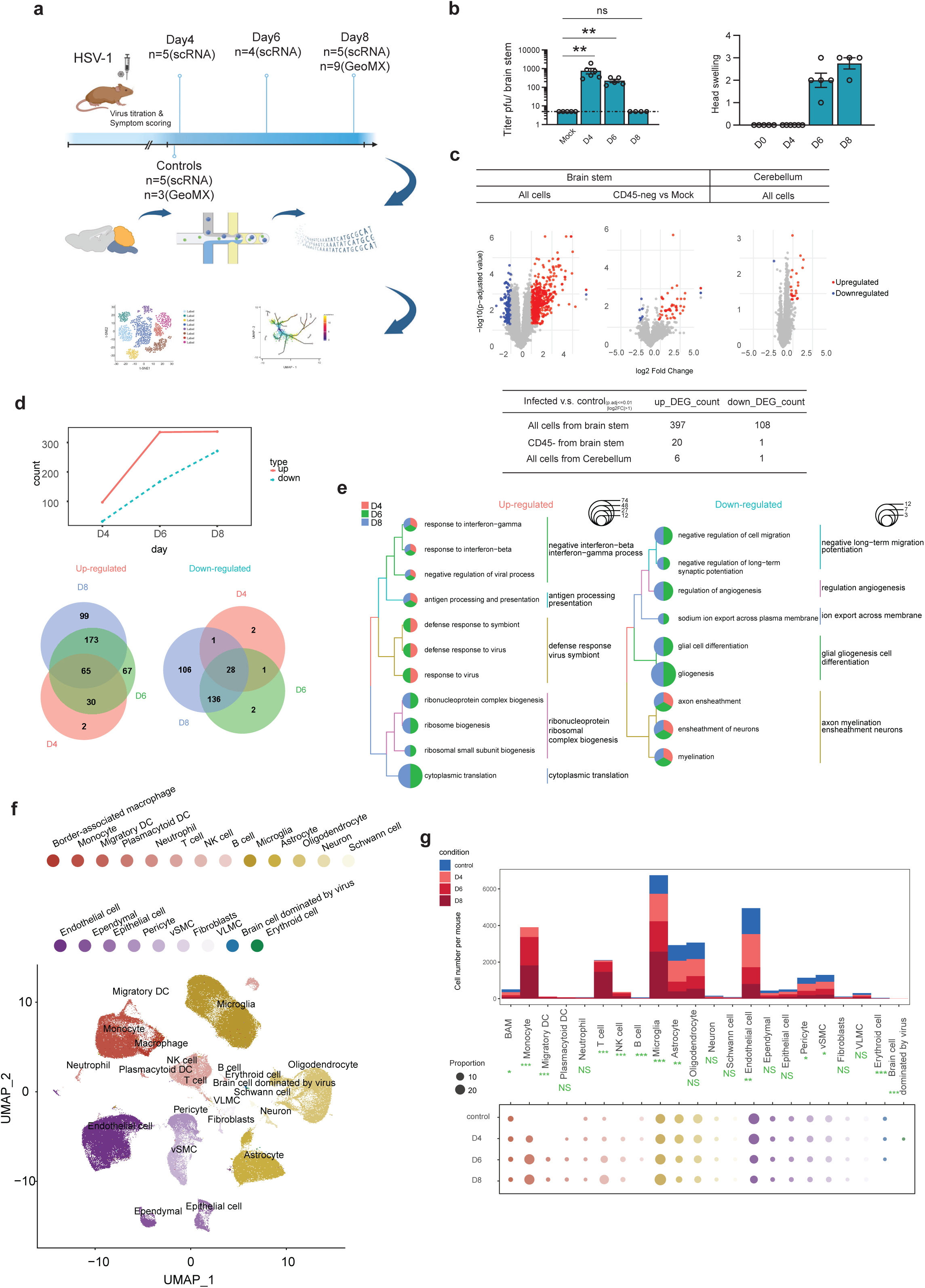
HSV-1 CNS infection leads to influx of immune cells and disturbance of composition and integrity of brain cells. **a,** Workflow for single-cell and spatial transcripts profiling of mouse brainstem cells. **b,** Bar plots illustrate the progression of viral titer and head swelling in the brainstem at various time points following viral infection. **c,** Volcano plots showing differential gene expression between HSV-1-infected and control mice in GeoMx spatial profiles. **d,** Number of differentially expressed genes in merged single-cell transcript profile from each time point after infection. Venn diagrams showing overlap of significant up-regulated and down-regulated genes between the time points. **e,** Comparison of enriched GO terms of up-regulated and down-regulated genes from different time point after infection. The size of each pie presents the number of genes affected. **f,** UMAP plot for dimensionality reduction and clustering from the sequenced mouse brain cells labeled with cell type annotation. **g,** Upper panel: cell type count number per mouse in control mice and at the three time points after HSV-1 infection. Lower panel: bubble plot of cell type proportion; dot color represent cell type identity and dot size represent proportion. Data were analysed using two-tailored one-way ANOVA. Asterisks (*) indicate p < 0.05, double asterisks (**) indicate p < 0.01, and triple asterisks (***) denote p < 0.001.

Based on the above information, we employed single-cell RNA sequencing to analyze changes in the transcriptome profile of the mouse brain stem on days 4, 6, and 8 following HSV-1 infections. A first broad analysis of the changes of the host transcriptome, showed that the fraction of reads mapping to antisense gene regions increased significantly on day 4 compared to the uninfected controls, followed by a decrease on subsequent days (Extended Data Fig. 1d), indicating general disturbance of transcription integrity, as seen previously in the context of HSV infections ^34^. Second, the mRNA transcripts from cells in the infected brain stems contained progressive sequence mismatch to the C57BL/6 genome, indicative of expressing of RNA-editing enzymes (Extended Data Fig. 1e). Third, we identified top correlated expressed host genes with viral transcripts at each time point (Extended Data Fig. 1f). Il1rn, encoding the IL-1 receptor antagonist emerged as the top gene correlating between viral and host transcripts in single cells, and we identified myeloid cells to be the main source of Il1rn.

To initially prioritize the most significantly differentially expressed genes (DEGs) at each infection time point, we aggregated the replicates of single-cell transcript profile within each time point into a consolidated transcriptome. Upon analysis of this dataset, we found that the number of up-regulated genes exceeded the number of down-regulated genes, and with a continuous increase in DEGs even beyond the time of presence of infectious virus (Fig. 1d). When exploring the hierarchical clustering of enriched function of DEGs the upregulated DEGs were found to be enriched for immune functions, including IFN responses, antiviral activity, and antigen presentation (Fig. 1e). Among the down-regulated DEGs, the enriched pathways included many functions related to physiological brain function, including gliogenesis, axon ensheathment and angiogenesis (Fig. 1e).

From the mice subjected to single cell RNA sequencing, a total of 143,915 high quality single cells were obtained. Our analysis revealed a comprehensive spectrum of 22 distinct cell types, encompassing 8 immune cell types, 5 brain resident cell types, 8 cell types involved in vasculature and tissue structure (Fig. 1f, Extended Data Fig. 1g). Following HSV-1 infection, influx of monocytes, dendritic cells (DC), and natural killer (NK) cells was detected from day 4, with particularly the number of monocytes increasing and continuing through the time course of the experiment (Fig. 1g). When looking at the resident brain cells and cells associated with vasculature and tissue structure, we observed that microglia numbers dramatically increased to become the most abundant cell type identified in the samples from mice infected for 8 days. By contrast, we observed a profound and temporally progressing decrease in the number of astrocytes in the HSV-1-infected brain stem. A similar loss of cells following infection was also observed for endothelial cells and pericyte, which are all the key constituents of the blood-brain barrier (BBB) (Fig. 1g, Extended Data Fig. 1h). This was not merely due to change in cell proportions, due to influx of immune cells, since analysis of the brain resident, vasculature, and tissue structure cells alone, also showed a reduction in astrocytes, endothelial cells, and pericytes (Extended Data Fig. 1i). Collectively, these data demonstrate that the HSV-1-infected brain is subject to a transcriptional stress response indicated by reduced transcriptional integrity, elevated immune activity, and down-modulation of homeostatic brain pathways. This is paralleled by profound increase in microglia, influx of monocytes, and loss of specific cell types, notably astrocytes, pericytes, and endothelial cells.

### Alterations in transcriptional profile following HSV-1 infection

Next, we analyzed the DEGs at finer resolution, with the aim to pinpoint comprehensive alterations in transcriptional profiles in the different cell types following viral infection. In line with the general observation that more DEGs were upregulated than down-regulated (Fig. 1d), this was also observed in most cell types (Fig. 2a). The microglia exhibited the highest number of DEGs, and upregulated genes included the chemokines *Ccl2*, *Ccl5*, and *Ccl12;* the IFN-stimulated genes (ISGs) *Ifi27l2a*, *Ifitm3*, and *Bst2*; and the MHC genes *H2-K1* and *H2Ab1* (Fig. 2b). Although of lower amplitude, most other cell types also showed early induction of many ISGs and chemokines, and later induction of antigen-presentation-related genes. This was confirmed in a pathway enrichment analysis (Fig. 2c, Extended Data Fig. 2a-f). Of note, the above analysis did not include monocytes, the most abundantly recruited immune cells, since they could not be compared to control brains, which did not contain monocytes.

**Fig. 2.**
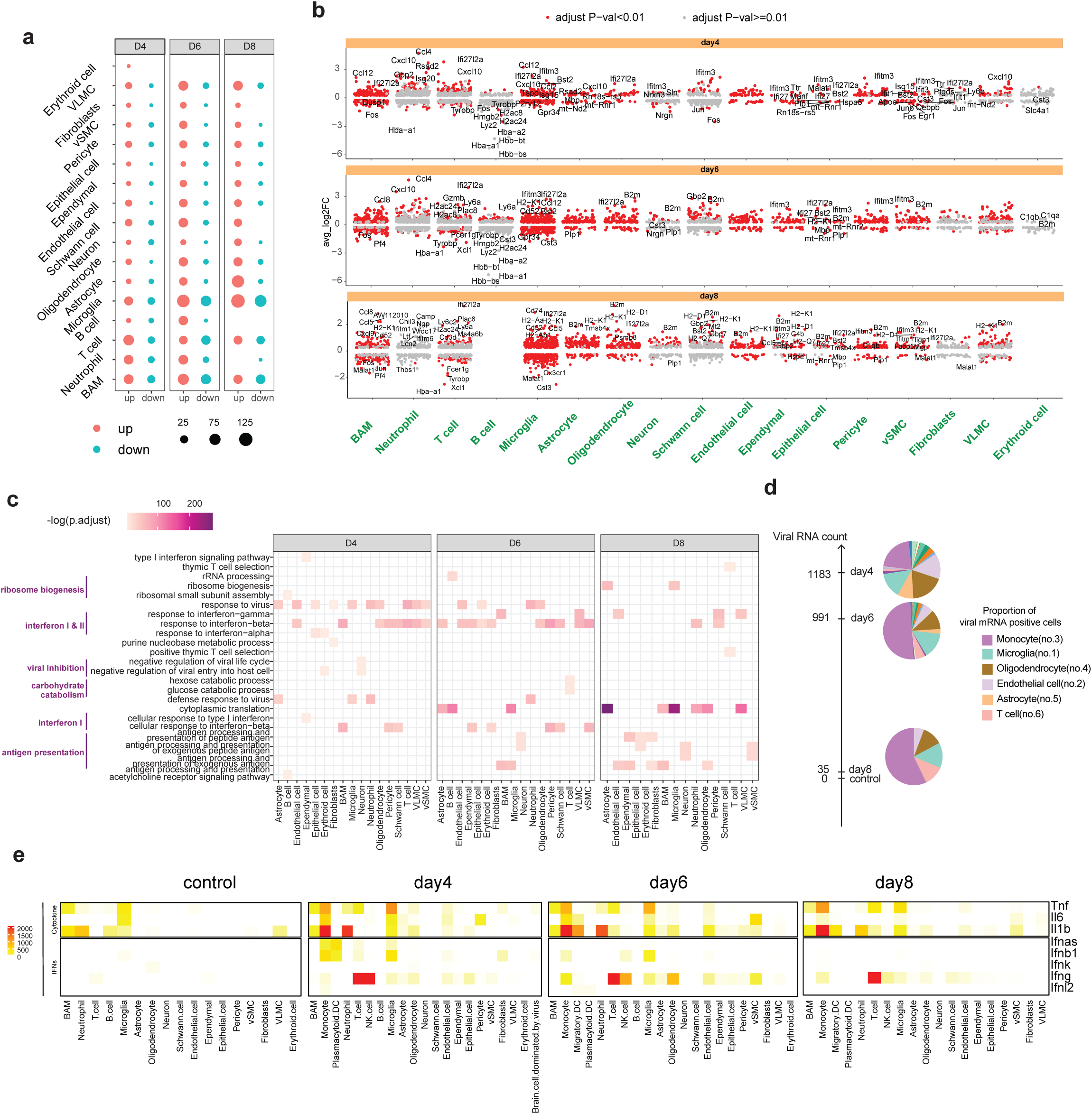
Alterations in transcriptional profile following HSV-1 infection. **a,** Bubble plot of number of differentially expressed gene in annotated cell type compared with control samples. **b,** Scatter plots of significantly differentially expressed genes shown in red for each cell type. Top genes labeled for all cell types. Differential expression testing is conducted using the non-parametric Wilcoxon rank-sum test. **c,** Enriched functions related to interferons, antiviral activity, antigen presentation, etc in each cell type across the different infection time points. **d,** Pie charts illustrating the distribution of viral transcript loads across various cell types on days 4, 6, and 8. **e**, Heatmap displaying the expression patterns of selected interferon and cytokine genes in the annotated cell type.

As a next part of the initial characterization, we first investigated the abundance of HSV-1 transcripts. Significant amounts of viral transcripts were observed on day 4 and 6 post infection (Fig. 2d), which was in line with the detection of infectious viral particles in the brain stem (Fig. 1b). Interesting, viral transcripts were found to be particularly enriched in monocytes and microglia (Fig. 2d), which have low permissiveness for HSV-1 infection ^4,34^, suggesting uptake of virus and virus-infected cells. Second, we wanted to investigate which cell types that expressed the cytokines playing apex roles in the immune cascades being activated in the HSV-1-infected brain. We noted that the type I IFN genes was mainly derived from microglia and monocytes, as well as the very low-abundant plasmacytoid dendritic cells, peaking on day 4, while a strong *Ifng* expression (type II IFN) was observed throughout the time of the infection from NK cells and T cells (Fig. 2e; Extended Data Fig. 2g). With respect to inflammatory cytokines, we observed strong induction of notably *Tnfa* and *Il1b* by monocytes and microglia, with monocytes being the main source and extending beyond the duration of viral presence on the brain (Fig. 2e). Collectively, these results show that the immune response in the HSV-1-infected brain is dominated by early IFN and antiviral responses and subsequent activation of antigen presentation, with most cell types being involved notably microglia and monocytes. Monocytes contribute substantially to both the type I IFN genes and prolonged expression of proinflammatory cytokines.

### Exploration of microglia activity and subpopulations

Given substantial effect of HSV-1 infection on the proportion and gene expression in microglia, we wanted to characterize this cell type in more details. Using Iba1 immunostaining in differences radial distances from the infection foci, zone 0-2 (Fig. 3a), we observed an absence of the characteristic ramified microglia in the center of the infection foci (zone 0, Fig. 3b), and an increase in the number of cells in the periphery and adjacent areas (zone 1 & 2 Fig. 3b-c). Comparing the morphological profile microglia showed a profound shift from the ramified surveillant shape abundant in the mock brains (type A, Fig. 3d) towards the hypertrophic and amoeboid shape (types C and D) at day 5 following infection, indicating activation (Fig. 3d). It is important to note that the amoeboid microglia cannot be distinguished from infiltrating monocytes through the method used. The amoeboid microglia/monocytes were more abundant in zone 1 than in zone 2 on day 5 (14% vs. 3%), while they were equally abundant in the two zones on day 8 (20% and 17%), where the virus had been cleared. In fact, at this point over 70% of the microglia in both zones showed the more activated profiles (type C and D) that were very rare in the homeostatic brain (Fig. 3b and d), suggesting an ongoing microglia response in areas distant from the infection foci.

**Fig. 3.**
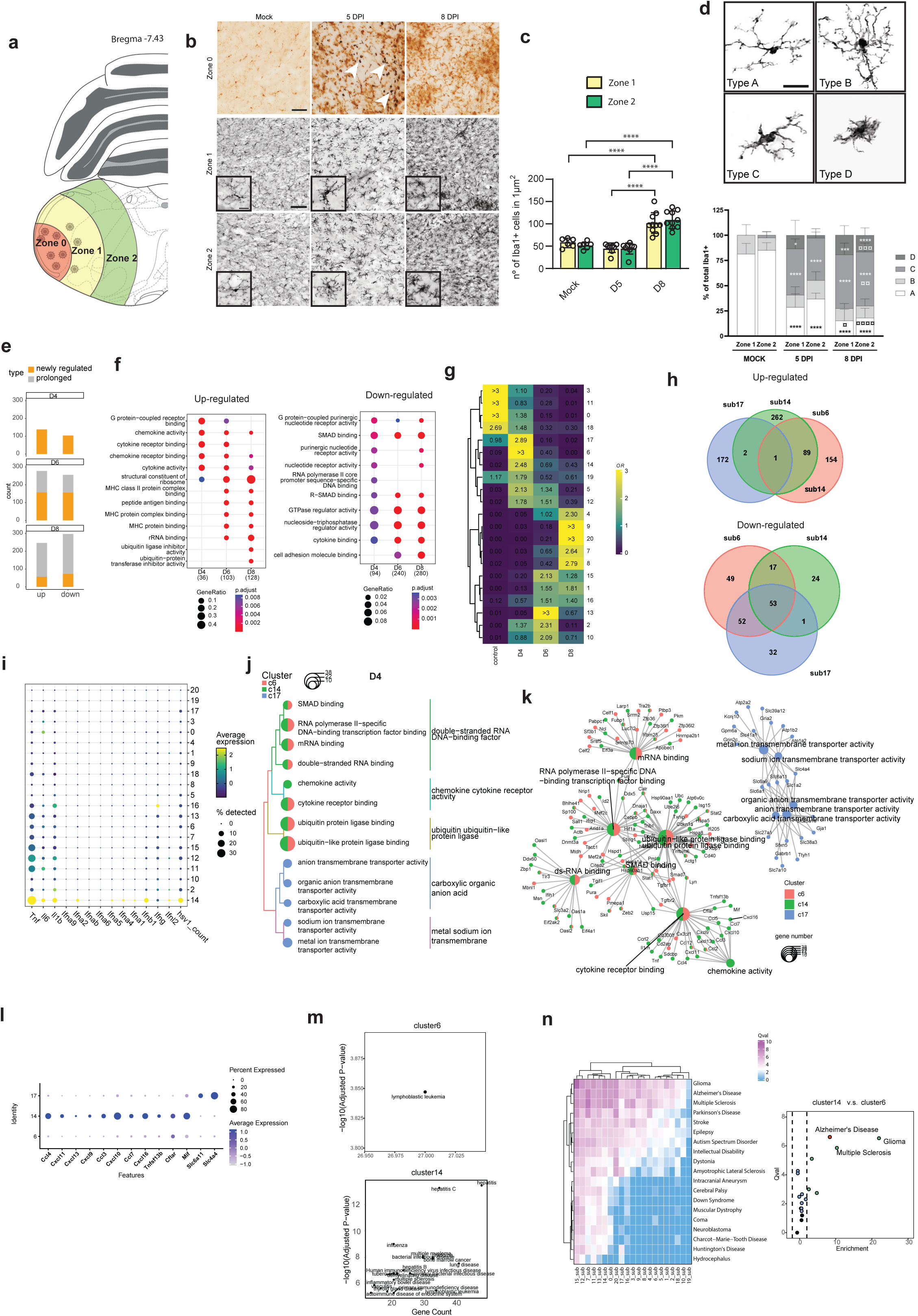
Transcriptional and functional heterogeneity within subpopulations of microglia. **a**, Illustration of area in the brain stem, categorized as zones 0, 1 (500-1000µm from center of virus focus), and 2 (1000-1500µm from center of virus focus). **b**, Representative images from double staining of brain stem sections with anti-HSV (black) and anti-Iba1+(brown). Representative HSV+ cells are shown by the white arrows. Scale bar, 50µm. Inserts display changes in the morphology of Iba1+ cells. Scalebar, 20µm. **c**, number of Iba1+ cells were quantified for all three conditions and in zone 1 and 2. **d**, Representative images of Iba1+ cells grouped in the different morphology categories A-D. Scale bar in 40x photos represents 50µm. Percentage of Iba1+ cells in the different morphology classes and in zone 1 and 2 were quantified. **e,** Bar plots showing the number of DEGs in microglia at various infection time points. Orange, newly regulated genes; gray, genes also differentially regulated at the previously examined time pointed (i.e. two days earlier). **f,** Comparison of enriched functional terms for upregulated and downregulated DEGs in microglia. **g,** Heatmap showing the prevalence of microglia subpopulations at the different time points, estimated by the Ro/e score. **h,** Venn diagram of DEGs in subpopulations #6, #14, and #17. **i,** Dot plot of selected cytokine and interferon genes, along with HSV transcripts. **j,** Comparison of enriched function term of up-regulated genes in subpopulations #6, #14 and #17. **k,** Network plot illustrating linkages between genes and biological concepts derived from enriched GO terms in subpopulations #6, #14, and #17. **l**, Highlight of selected genes within the enriched terms for microglia subpopulations #6, #14, and #17. **m**, Scatter plots of enriched human disease terms in subpopulations #6 and #14. **n,** The left panel presents a heatmap of brain disease-related pathway activity values in microglia subpopulations. The right panel compares subpopulations #6 and #14. Panel c, Data is presented as ±SD. Two-way ANOVA with matched values, followed by Tukey’s multiple comparisons test was applied (*<p=0.05 ****p≤0.0001). Mock, n=6 (left=3, right=3), 5 DPI, n=8 (left=4, right=4), 8 DPI, n=10 (left=5, right=5). Panel d: Data is presented as +SD. Two-way ANOVA with matched values when analyzing zone 1 and zone 2 within the same group, or without matched values when analyzing the same zone across groups. * compared to the same zone in the mock (*<p=0.05, ***p≤0.001, ****p≤0.0001). ¤ the same zone compared to 5 DPI (¤<p=0.05, ¤¤p≤0.01, ¤¤¤p≤0.001, ¤¤¤¤p≤0.0001). Mock, n=6 (left=3, right=3), 5 DPI, n=8 (left=4, right=4), 8 DPI, n=10 (left=5, right=5).

To explore changes in the transcriptomic profiles of microglia upon viral infection, we compared the overall transcriptomic alterations at different infection time points and found significantly greater changes in the microglial transcriptomes on days 6 and 8 compared to day 4 post-infection (Fig. 3e). Genes significantly upregulated in microglial cells on days 4 and 6 are predominantly associated with cytokine activity. Conversely, on days 6 and 8, significantly up regulated genes are enriched in functions related to antigen presentation and protein synthesis, the latter indicating cellular activity and proliferation (Fig. 3f left, Extended Data Fig. 3a). The genes downregulated in microglia were enriched in the pathways SMAD binding, cytokine binding, and cell adhesion molecules binding, which play critical roles in regulating microglial function, immune responses, and interactions within the CNS microenvironment (Fig. 3f right, Extended Data Fig. 3b). Alteration of these signaling pathways may promote changes in actin cytoskeleton organization and interactions with neighboring cells, leading to morphological shifts in microglia.

To investigate the heterogeneity of microglia following viral infection, we analyzed subpopulations by further dividing them into distinct subclusters. Condition enrichment scores reveal the distribution of subpopulations at the infection time points post infection (Fig. 3g). Importantly, there was a very dynamic change in the microglia population composition during HSV-1 brain infection. Further characterization of the subpopulations showed that the dominating subpopulations on day 4, #6, #14, and #17, were transcriptionally rather distinct, (Fig. 3h, Extended Data Fig. 3c). Most notably, subpopulation #14 was the one containing most HSV-1 RNA and expressing the highest levels of transcripts for type I IFN genes and the inflammatory cytokines *Tnfa, Il1b, and Il6* (Fig. 3i). This subpopulation also expressed the highest levels of chemokine transcripts, and high levels of cell-death-associated genes and nucleis-acid-binding proteins (Fig. 3j-l; Extended Data Fig. 3d). Importantly, subpopulation #14 was also associated with inflammation and disease phenotypes (Extended Data Fig. 3e), including viral diseases, and neurodegenerative diseases (Fig. 3m-n). Among the subpopulations that dominated on day 6 and 8, key activities were antigen presentation, proliferation and repair (Extended Data Fig. 3d). Subpopulations #15 and #16, which have high expression of *Top2a, Mki67*, and *Stmn1*, were classified as proliferating (Extended Data Fig. 3e), and both were enriched in functions related to tubulin binding (Extended Data Fig. 3f). These two subpopulations peak on day 6, with the former being distinguished by its enrichment in MHC protein binding function (Extended Data Fig. 3f). Finally, within the subgroups identified as enriched on day 8, subpopulation #9 stands out for its particularly high expression of Lgals3, a marker gene associated with clearance repair function (Extended Data Fig. 3e). In summary, HSV-1 infection leads to a profound change in the microglia phenotypes, and includes a highly antiviral and proinflammatory subpopulation, #14, which is present early and transiently in the HSV-1-infected brain microenvironment.

### Investigation of heterogeneity and differentiation within monocyte subpopulations

Recruited monocytes progressively accumulated in the infected mouse brain stems, reaching a proportion of over 20% by day 8. Within the large population of monocytes, 20 subclusters were identified exhibiting heterogeneous features revealed by distinct gene expression patterns of subsets (Fig. 4a, Extended Data Fig. 4a and 4b). Among the subpopulations abundant on day 4, particularly population #4 and #13, but also population #17, are characterized by ISG expression and antiviral responses (Fig. 4b, and 4c). Additionally, these subpopulations were enriched for regulation of vasculature (Fig. 4b), likely contributing to the loss of BBB integrity. Among these, subpopulation #4 contained the highest level of viral RNA and was categorized as Ly6c2 positive inflammatory monocytes with expression of all the cytokines *Tnf*, *Il1b*, and *Il6* (Fig. 4c, and 4d, Extended Data Fig. 4c). A small fraction of the cells from this subpopulation expressed high levels of type I IFN genes (Fig. 4d). In contrast to this, subpopulation #13 exclusively expressed high levels of *Il6*, and #17 predominantly expressed *Tnf* (Fig. 4d).

**Fig. 4.**
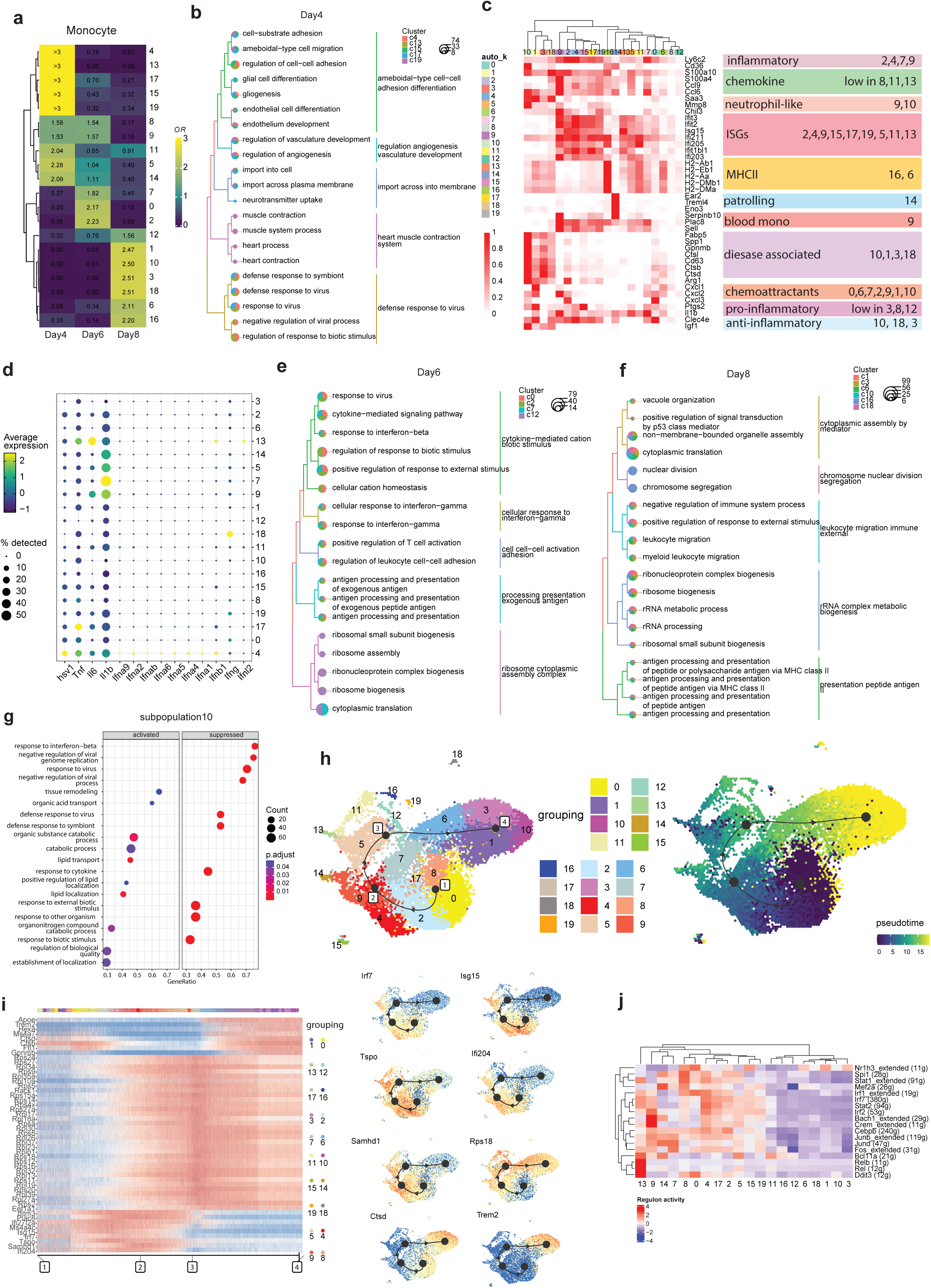
Heterogeneity monocyte subpopulations in the HSV-1-infected brain. **a,** Heatmap showing the prevalence of monocyte subpopulations at the different time points, estimated by the Ro/e score. **b,** Enriched function terms in up-regulated genes in subpopulations prevalent on day 4 post infection. **c,** Heatmap of characteristic marker genes for monocyte subpopulations, including functional annotation of the groups. **d,** Dot plot of viral transcripts and selected cytokine and interferon genes. **e,** Enriched function terms in up-regulated genes in subpopulations prevalent on day 6 post infection. **f,** Enriched function terms in up-regulated genes in subpopulations prevalent on day 8 post infection. **g,** Dot plots showing activated and suppressed terms from GSEA analysis in subpopulation 10. **h,** Left: UMAP of the inferred trajectory with connected milestone points, labeled by subpopulation identity. Right: UMAP of the inferred trajectory colored by pseudotime order. **i,** Heatmap overview of the most predictive genes along the inferred trajectory and selected feature plots. **j,** Heatmap showing the regulon activity scores of subpopulation-specific transcription factors.

As was also seen for the microglia subpopulations, the monocyte subpopulations showed high degree of dynamics, and the most abundant populations on day 4 post infection, were barely detectable on day 6 (Fig. 4a). The monocyte subpopulations dominating on day 6, #0, #2 and #7, were categorized as inflammatory and chemoattractant (Fig. 4c). The biological activities at this time point were characterized by a shift from pattern recognition receptor (PRR)-driven cytokine responses towards antigen-presentation and T cell activation, including IFN-γ responses (Fig. 4e). Among the monocyte subpopulations dominating on day 6, #7 stood out as being the main source of *Il1b* among all subpopulations (Fig. 4d). The transcriptome from the subpopulation showed enrichment of a range of proinflammatory activities, including regulation of leukocyte cell-cell adhesion and activation (Extended Data Fig. 4d). Interestingly, subpopulation #7 did not contain significant levels of viral RNA (Fig. 4d), thus suggesting a more complex mode of induction that PRR stimulation alone. In this regard, we noted that this subpopulation also showed enrichment in IFN-γ responses, known to be a major driver of monocyte/macrophage activation (Extended Data Fig. 4d). The strong down-regulation of processes associated with chemotaxis, and upregulation of processes involved in cel-cell adhesion suggests recent recruitment to the area (Extended Data Fig. 4d). Finally, the monocyte functions enriched on day 8 post infection still included antigen presentation, but also more regulatory activities of immune responses (Fig. 4f). Among the monocyte subpopulations identified on day 8, subpopulations #3, #10, and #18 expressed the C-type lectin Clec4e (MINCLE) (Fig. 4c). This protein has been reported to be a cellular sensor for necrosis ^35^, which is a signature of HSE ^36^, and may thus be central for clearance of debris from the HSV-1-infected brain. Additional functional enrichment analysis revealed that cluster #10 suppresses key inflammatory activities (Fig. 4g, Extended Data Fig. 4e). In contrast, subpopulations #6 and #16 notably expressed MHC-II related genes, thus suggesting that the pro- and anti-inflammatory activities of infiltrating monocytes is mediated by distinct populations of cells at this stage of the host response to the infection. The observation that subpopulations #16 is undergoing cell division (Fig. 4f), could indicate a prolonged presence in the post-infectious brain

Considering the prevalence at different time points and diverse antiviral functions, the differentiation trajectory of the recruited monocyte subpopulations was explored using comprehensive trajectory inference analysis (Fig. 4h, Extended Data Fig. 4f and g). The sequence of subpopulations appearing according to the pseudotime was consistent with the findings in real time (Extended Data Fig. 4h). The significant variability in gene expression along the trajectory also revealed a transition of recruited monocytes from an antiviral stage to a tissue homeostatic stage (Fig. 4i). The IFN-regulated genes *Ifi204*, *Irf7*, and *Isg15*, were all highly expressed at the onset of the immune response trajectory. This overlapped with, and was exceeded by, the expression of the activation marker *Tspo*, which in turn was followed by the proliferation marker *Rps18*. *Trem2* was highly expressed at the end stage of the recruited monocytes. *Trem2* plays a crucial role in regulating inflammation, phagocytosis, and the maintenance of tissue homeostasis. The inferred transcription factor regulon based on the single-cell transcriptional profile revealed more intensive regulon activity at the initial stage of the monocyte trajectory compared to later stages. Specifically, IRF7 and STAT2 regulate gene expression in subpopulation #4, NF-κB subunits Rel and RelB play significant roles in subpopulation 13, while Jun-Fos family members contribute to the activity of subpopulation #7 (Fig. 4j). Collectively, the monocytes recruited into the HSV-1-infected brain, show high functional diversity and dynamical change over time, enabling timely stimulating of innate and antiviral activities, coordinating of immune response, and resolving of inflammation.

### Transcriptional changes and cell-cell communication mediating disturbance of blood-brain barrier integrity

The early analysis of the scRNA transcriptomes revealed a significant reduction in cells associated with the BBB, i.e. astrocytes, pericytes, and endothelial cells (Extended Data Fig. 1h). Upon examining for access of Evans blue into the brain stem in HSV-1-infected mice, we observed that the exclusion of the dye in uninfected mice was impaired following infection, notable visible on day 6 and 8 post infection (Fig. 5a). Since the scRNA and GeoMx transcriptome data from the brain stem showed very strong correlation, and enrichment of the same GO terms (Fig. 5b and 5c), we reasoned that the scRNA sequencing data could be used to infer transcriptional changes and cell-cell communications in the BBB microenvironment. Focusing on endothelial cells, we found strong upregulation of IFN responses throughout the time span of the experiments, including both type I and II IFNs, and a later upregulation of processes supporting T cell responses (Fig. 5d). Importantly, however, at late times we observed downregulation of pathways associated with tissue homeostasis and integrity, as well as down-regulated of pathways involved in interaction between the neurovascular unit and neurons (Fig. 5e).

**Fig. 5.**
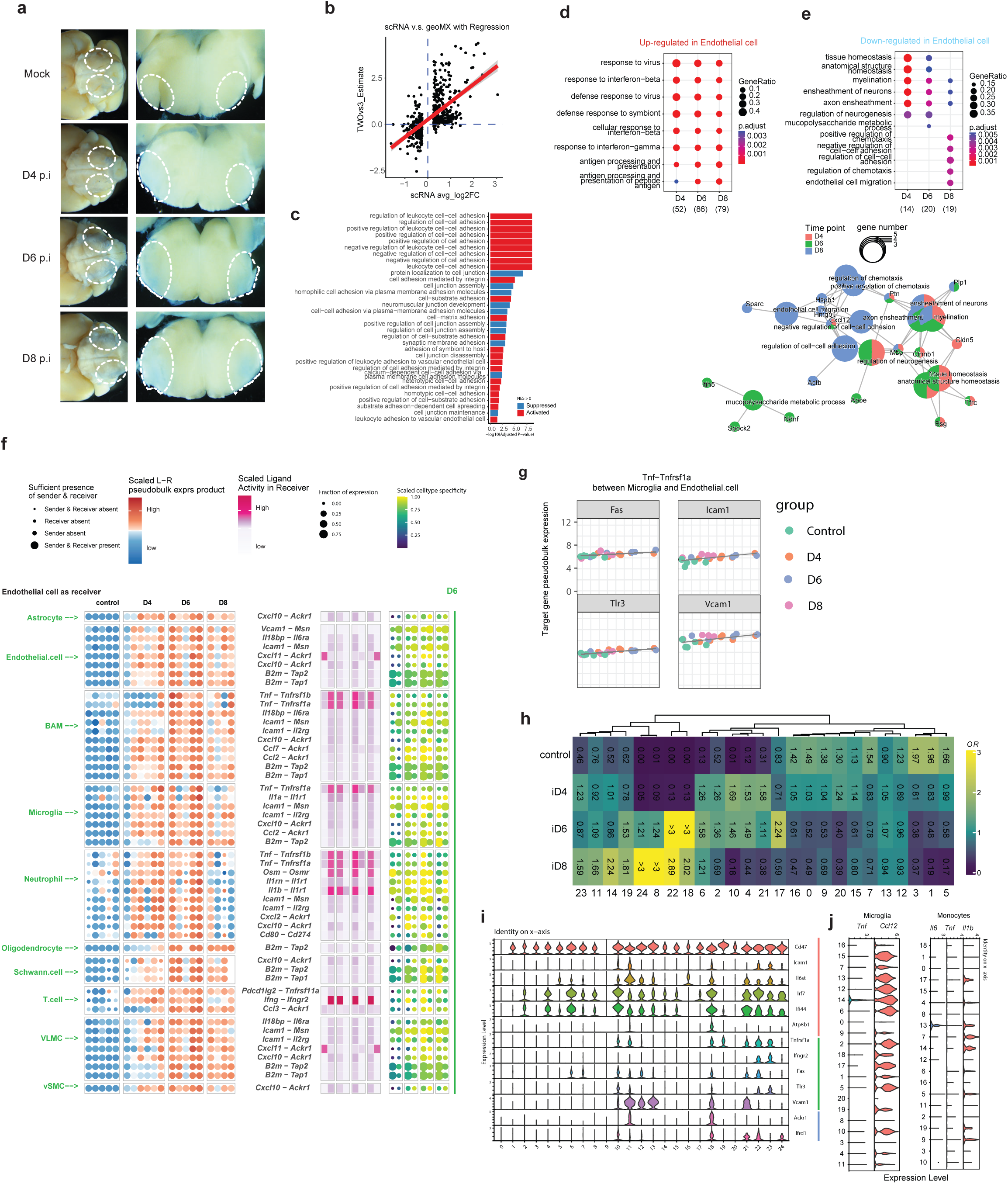
Transcriptional changes and dynamic communication in cells involved in the weakened blood-brain barrier post-infection. **a,** Mock- and HSV-1-infected mice received Evans blue dye by the i.p. route 4hr before transcardial PBS perfusion. Representative images of Evans blue dye leakage in mice brain stems from ventral and coronal positions are shown (5x and 2.5x objective, n=5 pr. group). **b,** Scatter plot showing the correlation between DEGs from merged single-cell transcriptome profiles and GeoMx spatial profiles on day 8 post-infection. **c,** Bar plots depicting activated and suppressed terms related to cellular adhesion and junctions from GSEA analysis in GeoMx spatial profiles. **d,** Comparison of enriched function term of up-regulated genes in endothelial cell. **e,** Upper part: Comparison of enriched function term of down-regulated genes in endothelial cell. Lower part: network of linkages between genes and biological concepts from enriched down-regulated DEGs. **f,** Plots of receptor and ligand communication pairs differentially expressed under various conditions: the red and blue dot plots represent per-sample pseudobulk expression of communication pairs, the heatmap displays scaled ligand activities, and the yellow and green dot plots indicate cell type specificity. **g,** Scatter plot shows the pseudobulk expression of Tnf-Tnfrs1a pairs and the expression of target genes between microglia (as senders) and endothelial cells (as receivers). **h,** Heatmap showing the condition prevalence of each endothelial cell subpopulations, estimated by the Ro/e score. **i,** Selected receiver genes expression in subpopulations of endothelial cell. **j,** Selected ligand genes expression in subpopulations of microglia and monocytes.

Next we used NicheNet to infer what cell-cell communications that could be responsible for the observed changes. First, we noted that the cross-talk between cells forming the BBB were impaired in the infected brain stem (Fig. 5f, Extended Data Fig. 5a). This included for instance the interactions between astrocytes and endothelial cells through Vascular Endothelial Growth Factor A (VEGFA) and its receptors as well as between pericytes and endothelial cells through Transforming growth factor beta (TGFβ) and Angiopoietins (Extended Data Fig. 5a). Second, we found that the inflammation evoked by the infection led to several new cell-cell interactions for endothelial cells known to promote break-down of the BBB. Most notably, the *Tnf*, mainly expressed by microglia and monocytes, but also other cell types (Fig. 1e), extensively stimulated endothelial calls (Fig. 5f-g, Extended Data Fig. 5a-b). Additionally, *Ifng* expression by T cells was accompanied by induction of IFN-γ-induced gene profiles in endothelial cells (Fig. 5f, Extended Data Fig. 5a). These results identify pathways that likely contribute to disturbance of the BBB during HSV-1 CNS infection.

Finally, we subdivided the endothelial cells into subpopulations and identified 24 transcriptionally distinct clusters (Extended Data Fig. 5c). In line with the observed break-down of BBB integrity, the major changes in subpopulation composition occurred on day 6 and 8 post infection, with emergence of populations #8, #22, and #24, and to a lesser extent population #14 and #18 (Fig. 5h). Several of these populations were categorized to disease- or damage-associated phenotypes (Extended Data Fig. 5d), although they generally did not contain high level of viral transcripts (Extended Data Fig. 5e). This suggests that the damage to the BBB is caused by inflammation rather than viral. In line with the disease- or damage-associated phenotypes, populations #18, #22, and #24 showed high expression of several genes involved in pyrotosis and apoptosis (Extended Data Fig. 5f). This was also seen in astrocytes, which were not examined in details, but expressed high levels of *Gsdme*, encoding a death executer in pyropsosis (Extended Data Fig. 5g). Subpopulation #14 exhibited a proliferative transcriptome profile, potentially indicative of initiation of repair activities. Finally, when looking into the cell-cell cross-talk between subpopulations in the different cell types, we found that IFN-γ and TNFα receptors *Ifngr2* and *Tnfrsf1a* were highly expressed in the endothelial subpopulations suspected to be involved in BBB damage, including #18 and #22 (Fig. 5i).

These cells received cytokine signals from microglia subpopulation #14 and multiple monocyte subpopulations, including #14, #17, and #7 (Fig. 5j). Collectively, these data show that HSV-1 infection in the CNS evokes an inflammatory response, which disturbs the steady-state activity of the brain vasculature and breakdown of the BBB, thus facilitating influx of immune cells in the brain, but also allowing influx of unwanted molecules and alterations of homeostatic interactions in the neurovascular unit.

## Discussion

Viral encephalitis is a devastating disease, and can have fatal outcome, or lead to severe morbidity ^1^. The immune system is essential for control of viral CNS infections, but can also contribute to disease development ^37,38^. In this work, we show that HSV-1 infection in the CNS leads to a profound transcriptional change in many cell types, recruitment of a panel of immune cells, which drive both protective and pathological responses. In particular we identify specific microglia and monocyte subpopulations, and uncover their phenotypic profiles and interaction networks, and suggest how they contribute to the outcome of infection. This includes both control of infection and disturbance of homeostatic functions, most notably the integrity of the BBB.

An initial overall characterization of the transcriptional changes during the course of HSV-1 brain infection showed strong upregulation of immune-related genes, and most DEGs where indeed expressed by CD45+ cells, i.e. immune cells. The dominant cell types were in addition to microglia, monocytes, and later T cells, all of which continued to increase after the infection was cleared, in agreement with previous reports ^39^. At the single-gene level we observed the strongest correlation between viral transcripts and levels of *Il1rn* in single cells. Interestingly, this is in agreement with clinical data showing correlation between *IL1RA* and HSE disease ^40^, thus providing support for the translational potential of our results. We also found the transcriptome of the infected brain to contained elevated levels of antisense transcripts, and mismatch with the genome sequence. This is indicative of a stress-related transcriptional response in infected areas ^41,42^. Finally, we observed downregulation of a series of pathways involved in homeostatic brain activity. The latter is consistent with the clinical observation that the patients recovering from encephalitis may suffer several long-term effects, such as cognitive defects ^43^.

Microglia is the main brain-resident immune cell type, and is well-described to express most pattern recognition receptors and proinflammatory cytokine receptors ^44^. Previous studies have shown that microglia are essential for host defense against viral CNS infections with both RNA and DNA viruses ^10,45,46^. Depletion of microglia was found to decreased expression of IFNβ and impaired activation of T cells ^10,45,46^. In our study we observed increased abundance of microglia in the brain stem following HSV-1 infection, and also that microglia are a prime source of transcripts for *Ifnb1* and *Tnf*, and to a lesser extent also *Il1b* and *Il6*. Interestingly, we observed that microglia undergo a profound and diverse change in subpopulation composition upon infection, with none of the subpopulations being present in the uninfected brain stem areas sequenced being detectable on day 6 post infection. Generally, we observed a dynamic change from homeostatic subpopulations, to early interferonic/inflammatory subpopulations in the presence of virus, and eventually to proliferative and antigen-presenting subpopulations at late time points. Notably, a microglia subpopulation, #14, was highly abundant on day 4 p.i., expressed the highest levels of type I IFN and chemokine genes as well as *Tnf*. This population showed the highest abundance for viral RNA among the microglia subpopulations, suggesting this subpopulation to be the key virus-sensing cell population, orchestrating IFN-mediated antiviral defense and recruitment of cells from the periphery. Microglia #14 was involved in numerous interactions, including endothelial cells, thus indicating involvement in both disruption of the BBB and activation of infiltrating immune cells. Given the importance of microglia an antiviral defense ^10,45,46^. Future studies should explore whether the identified subpopulation #14 is specific for HSV-1 infection, or in fact represent the virus-activated microglia, similar to the inflammation-associated microglia emerging following lipopolysaccharide exposure ^47^, and the disease-associated microglia found in neurodegenerative diseases ^48^.

Monocytes are not present in the brain during steady state, but are recruited in high numbers to the brain in response infections with HSV-1 and other viruses ^39,49,50^, as well as non-viral acute brain condition, like trauma ^51^. We found that the recruited monocytes underwent a highly dynamic change, illustrated by rapid change in the composition of subpopulations. At the early time-points, the monocytes were characterized by antiviral activity, but also alteration of the endothelium and glial cell activity. At later time points this shifted to cross-talk with T cells, including responsiveness to IFN-γ and antigen-presentation, as well as proliferation and regulatory processes. Regarding cell-cell cross-talk we found that subpopulations #13, #17, and #7 among others interacted with endothelial cells through the multiple pathways, including TNF, IL-6, and IL1β may thus contribute to the damage to the BBB. Similar to microglia, the monocyte population that contained the most viral RNA, #4, also produced the highest levels of type I IFN genes. However, unlike microglia, expression of inflammatory cytokines did not correlate with levels of viral transcripts and was high in many subpopulations. This could indicate that the inflammatory response in monocytes is driven by more complex mechanisms than merely viral sensing by PRRs, possibly involving cytokine stimulation, metabolic alterations, and epigenetic alterations. In agreement with this, we observed monocytes to represent a more prolonged source of inflammatory cytokines beyond the duration of infection, and may therefore contribute to post-infectious brain damage. The expression of inflammatory cytokines by monocytes not containing high amounts of viral RNA, could also allow for monocyte-driven inflammatory activities in brain areas not directly infected by HSV-1. Of note, recent data based on HSV-1 infection in human brain organoids have identified TNFα as a key contributor damage of neuronal processes and neuroepithelium ^28^. The role of recruited monocytes to antiviral defense and brain pathogenesis needs further investigation.

Disruption of the BBB is a hallmark of many severe brain diseases, including viral encephalitis ^52,53^. Breakdown of the BBB is associated with numerous pathological processes, including access of large biomolecules from the blood stream into the CNS, disturbance of the mechanisms ensuring delivery of oxygen and nutrients into the brain, facilitation of influx of immune cells into the CNS, and can increased permeability of blood vessels leading to vasogenic edema ^52^. HSE is associated with brain edema, and we have previously reported this mice, occuring in the infected part of the brainstem ^4^. We found that the brains of HSV-1-infected mice were permissive to Evans blue, and also that there was an unproportionally large loss of BBB-associated cell types in the infected brains. Previous studies have reported that brain infections with JEV, Rabies virus, coronavirus, Reovirus, and WNV also leads to breakdown of the BBB ^29,54–57^. For most of these infection models, IFNγ contributed to the disruption of the BBB through down-regulation of tight junctions ^54–57^, whereas in the WNV model, it was dependent on TNFα signaling ^29^. Yet other studies have implicated a role for IL1β in BBB breakdown, including activation of the endothelium and recruitment of activated proinflammatory leukocytes ^58,59^. Our observation of loss of BBB-associated cells is still not fully explained mechanistically and may well also include different forms of inflammatory cell death. In fact, we observed high expression of a range of genes involved in pyroptosis and apoptosis in the endothelial cell populations emerging late during infected and exhibiting disease/damage-associated transcriptome profiles. Based on the low abundance of viral RNA in BBB-associated cells, we do not favor the hypothesis that the loss is caused by lytic virus replication. Thus, the disruption of BBB integrity in the HSV-1-infected brain may include several mechanisms. First, rapid and prolonged downregulation of cross-talk that generate and maintain the BBB, including TGFβ, VEGFA and Angiopoietin signaling, from different cell types towards endothelial cells. Second, down-regulation of a panel of tight-junction-related genes. Third, elevated inflammatory signaling to endothelial cells, notably IFN-γ signaling from T cells and TNFα, IL1b, and IL-6 signaling from specific monocytes and microglia subpopulations, may promote disruption of barrier integrity. Fourth, adhesion of activated proinflammatory leukocytes, which produce e.g. inflammatory cytokines, reactive oxygen species, or matrix metalloproteases. Fifth, loss of BBB-associated cell types, likely involving different forms of programmed cell death. The points listed above may well be mechanistically connected, which will have to be experimentally tested in future work aiming to uncover the exact mechanisms governing disruption of the BBB during HSV-1 brain infection.

In this work, we report a temporally resolved overview of the single cell transcriptome during HSV-1 infection in the CNS. Through this analysis we identify a microglia subpopulation that is likely a central mediator of the early antiviral response, and recruitment of immune cells from the periphery. We also identify activities of one recruited cell population, monocytes, and highlight their role in both host defense, but potentially also in pathological inflammation. We illustrate this with the cross-talk between specific immune cell populations and endothelial cells associated with disruption of BBB integrity. One important lesson from this study is that host responses change very dynamically after acute challenge, hence highlighting that scientists should be careful with interpretation of data from one time point when studying acute diseases. HSE is a devastating disease, and the underlying mechanisms remain poorly described. Understanding of the mechanisms and dynamics in cellular transcriptomes, activities, and interaction patterns in the HSV-1-infected brain may lead to new insight into HSE, and lead to novel therapeutic opportunities.

## Methods

### Mice

C57BL/6 mice were bred at Taconic M&B (immunohistochemistry), and Janiver-Labs (single-cell sequencing). Isoflourane (Abbott), or (Ketamine (MSD Animal Health) and Xylazin (RompunVet)) was used to anaesthetize mice. Prior to experiments, mice were kept at the animal facility Faculty of Health Science, University of Aarhus. All mice used in this study were age-matched (7–12 weeks of age) male mice. All efforts were made to minimize suffering and mice were monitored daily during infection.

### Viruses and reagents

Dulbecco’s Modified Eagle Medium (DMEM) was obtained from BioWhittaker and supplemented with 100 IU ml^-1^ penicillin, 100 mg ml^-1^ streptomycin and LPS-free FCS (BioWhittaker). HSV-1 (strains McKrae) was grown in Vero cells. The Vero cells used were from the lab stock. The titres of the stocks used were 5-10X10^9^ PFU ml ^-1^. Titres were determined by plaque assay on Vero cells. Beriglobin was used to neutralize extracellular HSV-1 (CSL Behring) and fully neutralized the virus in the dilutions used to calculate titres.

### Murine ocular HSV-1 infection model

Mice were anaesthetized with intraperitoneal (i.p.) injection of Ketamine (100mg kg^-1^ body weight) and xylazine (10mg kg ^-1^ body weight). Corneas were scarified in a 10X10 crosshatch pattern and mice were either inoculated with 2X10^6^ PFU HSV-1 in 5ul of infection medium (DMEM containing 200IU ml ^-1^ penicillin and 200ug ml ^-1^ streptomycin), or mock infected with 5ul of infection medium. Mice were killed at the specified times post infection. Daily monitoring of weight and disease symptoms was performed during the infection period. Disease scores were determined as follows: head swelling (0, no bump; 1, minor bump; 2, moderate bump; 3, large bump. Mice that reached the humane endpoints or weight loss equal to or higher than 20% of initial weight were euthanized.

### Virus plaque assay

Brain stem was homogenized in a 2 mL Eppendorf tube by adding 500mL of Dulbecco’s phosphate-buffered saline (DPBS) (Sigma-Aldrich) and a stainless-steel metallic bead (Qiagen) per sample in Tissuelyser (II) (Qiagen) for 5 min at a frequency of 50 Hz. The homogenized isolated tissue was centrifuged and the supernatant was used for the plaque assay. Vero cells seeded in 6-well plates were inoculated with 100mL of serially diluted samples and 900mL of DMEM with 2% FBS. After 1 h of incubation, DMEM with 2% FBS and 0.4% human immunoglobulin was added to each well to neutralize any extracellular HSV-1. The plates were subsequently incubated for 2 to 3 days until virus plaques were visible and not overlapping. Staining with 0.03% methylene blue (laboratory stock) was used to visualize and allow quantification of the plaques.

### GeoMx

#### Tissue Specimens

The mice were perfused, and the dissected brains were fixed with 4% formaldehyde, and then embedded in paraffin blocks. Tissue microarrays (TMA) were made from regions of interest within the brain stem and cerebellum, and by extracting 2mm tissue cores from the paraffin blocks, followed by re-embedding these specimens into a single TMA block consisting of 43 cores in total. The GeoMx DSP has the constraint of only being able to scan a slide area of 36,6 x 14,6 mm, so the TMA block was constructed such that the cores were confined to this area. Sections of 5µm thickness were obtained from the TMA and were positioned in the center of the slide.

#### Tissue Preparation

The mounted sections were processed according to the manufacturer’s protocol (Nanostring Technologies MAN-10150-02). Sections were baked at 37°C overnight followed by 60°C for 1 hour. Deparaffinization was performed by incubation in Xylene (VWR cat. nr. 28975.360) three times for 5 min. each. Rehydration was performed with 100% ethanol (EtOH) twice for 5 min. each, 95% EtOH once for 5 min., 70% EtOH once for 5 min., and 1X phosphate buffered saline (PBS) (BioNordika cat. nr. BN-53100) once for 1 minute. Antigen retrieval was performed by submerging the slide into a 1X Tris-EDTA solution (Invitrogen, eBioscience cat. nr. 00-4956-58), at 80°C for 15 min., followed by a wash in 1X PBS for 5 minutes. Target retrieval was performed by submerging the slide into a solution containing 1X PBS and 0,1 µg/mL proteinase K (Invitrogen cat. nr. AM2546) at 37°C for 15 min., followed by a wash in 1X PBS for 5 minutes. Sections were postfixed in 10% neutral buffered formalin (NBF) (Sigma-Alrich cat. nr. HT501128), followed by two washes in a NBF stop buffer solution containing 2,42 g Tris base (Sigma-Aldrich cat. nr. T1503), 1,5 g Glycine (Sigma-Aldrich cat. nr. G7126), and 200 mL nuclease-free water (BioNordika cat. nr. BN-51100), for 5 min. each, followed by a wash in 1X PBS for 5 minutes.

Hybridization of photocleavable DSP RNA detection probes was performed by applying 220 µL hybridization solution to the tissue sections. The hybridization solution contained 25 µL DSP RNA whole transcriptome detection probes (GeoMx NGS RNA WTA Ms, Nanostring Technologies cat. nr. 121401103), 25 µL nuclease free-water, and 200 µL Buffer R (GeoMx RNA Slide Prep Kit for FFPE, Nanostring Technologies cat. nr. 121300313). The slide was then covered with a hybridisation coverslip and incubated at 37°C overnight. The following day the slide was dipped into a 2X saline sodium citrate solution (SSC) (Rockland Immunochemicals cat. nr. ROCKMB-045) to remove the coverslip, followed by two washes in stringent wash solution containing 4X SSC and 100% formamide (APPLICHEM cat. nr. 11237117), at 37°C for 25 min. each, followed by two washes in 2X SSC for 2 min. each. Sections were then blocked with Buffer W (GeoMx RNA Slide Prep Kit for FFPE, Nanostring Technologies cat. nr. 121300313) and incubated at room temperature for 30 minutes. The TMA slide was stained with fluorescent labelled primary antibodies for CD45-APC (BioLegend 30-F11, cat. nr. 103112) for 2 hours. The antibodies were diluted in Buffer W at 1:20. After incubation the slide was washed in 2X SSC twice for 5 min. each. DNA was counterstained with 400 nM SYTO 83 (Thermo Fischer cat. nr. S11364) and incubated at room temperature for 1 hour, followed by two washes in 2X SSC for 5 min. each. The slide was stored overnight in 2X SCC at 4°C.

#### ROI Selection Strategy

The slide was loaded onto the GeoMx DSP instrument and scanned to produce a digital image of the tissue sections. The digital image was used to visualize tissue morphology based on the fluorescent labelled antibodies CD45, and the DNA stain SYTO 83. Regions of interest (ROI) were selected based on high morphological expression of immune-cell rich areas with a high expression of CD45, and SYTO 83 and expression of HSV-1 staining on a sequential section, Multiple ROIs of various sizes (up to maximum ROI size of 660 x 785 µm) and containing varying cell counts were selected.

#### ROI Segmentation/AOI Profiling and AOI Collection

After ROIs were selected, the image analysis software integrated in the GeoMx DSP instrument was used to carry out threshold-based segmentation, to further divide each ROI into different segments. Segmentation into individual biological compartments was based on tissue morphology, CD45 expression, and cell count. Each ROI was divided into two segments: a segment containing individual cell populations with high CD45 expression, and a second segment containing cells with a positive nuclei stain but negative for CD45. Each segment within a ROI corresponds to one area of illumination (AOI). After segmentation of all ROIs into segments/AOIs, each AOI was exposed to ultraviolet (UV) light from the programmable digital micromirror device (DMD) built into the GeoMx DSP instrument. Each AOI was exposed to UV light in order to cleave DSP barcode tags from the AOI in question, releasing the tags from the tissue section. Released tags were collected by an automated microcapillary and dispensed into a 96 well microplate. After collection of all AOIs the microplate was sealed and stored at -20°C.

#### Library Preparation and Sequencing

Library preparation was carried out according to the manufacturer’s protocol (Nanostring Technologies MAN-10153-03). The collected DSP tag aspirate was thawed and dried on a thermocycler at 65°C, followed by resuspension in 10 µL nuclease-free water. A 4 µL aliquot of each resuspended aspirate was amplified in a PCR reaction containing NanoString SeqCode primers and PCR Master Mix (GeoMx Seq Code Pack_EF, NanoString cat. nr. 121400203). PCR products were pooled, and then purified with two rounds of AMPure XP beads (Beckman Coulter cat. nr. A63882). Libraries were quality controlled using a 4200 Tapestation (Agilent) and QuBit v2 (Fischer Scientific), before being sequenced on an NovaSeq 6000 (Illumina).

#### Data Analysis

The generated FASTQ files were converted to output DCC files using the GeoMx NGS Pipeline v2.5.1. In brief, raw reads were trimmed to remove adapter sequences. Overlapping paired-end reads were merged, and the resulting stitched reads were aligned to barcodes in the reference assay, hence creating aligned reads. The aligned reads were used to assign raw counts to biological target names. Reads matching the same barcode (PCR duplicates), were removed using the UMI region of each read, resulting in deduplicated reads converted to digital counts. Probe level counts were converted to gene level counts with the R package GeomxTools (v3.0.1). A limit of quantification (LOQ) for each segment was calculated as the geometric mean times the square of the geometric standard deviation of a set of negative probes. This LOQ was used as a segment specific gene filtering threshold, so that genes below LOQ in more than 2% of segments were filtered. Gene expression was subsequently upper quantile normalized. Differentially expressed genes were calculated using linear mixed models (package lmer4 v1.1) with mouse individual as a random factor. The groups (CD45-neg cells in infected mice versus all cells Mock ROI) and (all cells in infected and all cells in mock ROI) were compared. The biostatistical analysis of the differential expression matrices were conducted using the R Project for Statistical Computing(v4.3.3) and computed on GenomeDK. A gene set enrichment analysis (GSEA) were performed using clusterProfiler(v4.10.0) by applying the genome wide annotation for mouse(v3.18.0). For statistical significance of GSEA results a cutoff value of <0.05 were used for observed enrichments of gene sets. Enriched terms of interest were selected for further analysis using gene-concept network visualization to identify active genes, their relative regulation, and statistical significance of regulation using DOSE(v3.28.1).

### Evaluation of BBB Permeability

A 2% solution of Evans Blue in normal saline (4 mL/kg of body weight) was injected intraperitoneally. The stain was allowed to circulate for 4 hours. Afterwards, the mice were then transcardially perfused with 25 ml of PBS and fixed in 4% formaldehyde. The brainstems were dissected into 2 mm coronal sections. Evans blue dye leakage in mice brains from coronal and ventral positions were imaged on Leica M165FC microscope.

### Tissue immunofluorescent staining and immunohistochemistry

The brains were sectioned in the posterior-anterior direction on a freezing microtome (Brock and Michelsen, Thermo Fisher Scientific) into 40µm thick coronal sections and separated into a series of 10. Sections were stored at -20°C in an anti-freeze solution until use.

Immunohistochemistry was performed on free-floating sections. During the staining process, sections were washed multiple times in potassium-phosphate buffer (KPBS) between each incubation period. All incubation periods contained 0.25% Triton X-100 in KPBS. The sections were quenched for 10 minutes in a solution of 3% hydrogen peroxide and 10% methanol, followed by one hour of blocking with the appropriate 5% serum. The sections were incubated overnight at room temperature with the first primary antibodys monoclonal anti–HSV-1 (1/500, VP5 Abcam, clone 3B6), anti-mouse CD3ε Antibody (1/500 Biolegend, clone 145-2C11) or monoclonal rat anti mouse CD68 (1/500 Bio-Rad, clone FA-11) in 2.5% serum. The next day, sections were pre-blocked for 10 min in 1% serum before 2 hours of incubation with the biotinylated secondary antibody in 1% serum, (1/200, anti-mouse, pre-absorption performed in mouse tissue before adding to experimental tissue), followed by 1-hour incubation with avidin-biotin-peroxidase complex (Vectorstain, ABCkit, Vector Laboratories) in KPBS. Development was done with 3,3-diaminobenzidine (DAB) and 0.04% Nickel.

After the development of the first staining for HSV-1, the sections were washed and the above-described processes were repeated on the same sections for staining of Iba1+ cells (1:700, Wako Fujifilm, 019-19741). Iba1 staining was developed with DAB. Sections were mounted on gelatin-coated glass slides and coverslipped.

### Quantification of Iba1+ cells in tissue sections

Two sections containing the virus were identified per animal (mock, n=3, 5 DPI, n=4, 8 DPI, n=5) (Section 1: bregma -7.48 to bregma -7.76, section 2: bregma -7.20 to bregma -6.84). Each hemisphere of the sections was divided into three areas starting from the edged of the tissue: 0-500µm, which contained the most virus, 500-1000µm, and 1000-1500µm away from the area containing the most virus. In each of the 500-1000µm and 1000-1500µm areas, two photos (0.5mm x 0.5mm) were taken per section in both hemispheres. Extended focus imaging (EFI) photos were taken using an Olympus VS120 upright widefield slice scanner and the 40x objective. The z-range was set to 30µm and the z-spacing to 1 µm.

Each photo was analyzed in Fiji. Images were converted to 8-bit, and brightness/contrast was adjusted by the auto-function. The cell nuclei were counted using the multi-tool. The threshold was adjusted using the default setting. The analyze particle function (size: 0-infinity, circularity: 0-1, overlay masks) was used to obtain the %area covered by positive staining. The Region of interest (ROI) function was used to exclude any artifacts or vessels in the tissue. The average % Area of 500-1000µm and 1000-1500µm was calculated for the left and right hemispheres. The infection was not equally effective in both sides of the brain and each animal is represented by two data points: one for the left and one for the right hemisphere. A two-way ANOVA with matched values followed by Tukey’s multiple comparison test was applied.

The microglia morphology was evaluted in the previously described acquired photos. This analysis was performed in section 1 of each animal. Two photos from both zone 1 and zone 2 from both hemispheres were analyzed. The morphology of Iba1+ cells was determined by the length, thickness, number of processes, and characteristic of the cell body. Four profiles were defined: Type A (surveillant); no visible cytoplasm, small round dense nucleus, and long thin processes with little secondary branching. Type B (hyper-ramified); visible cytoplasm, round dense nucleus, and long processes with secondary branches. Type C (hypertrophic): Elongated and irregular cytoplasm, enlarged and less defined nucleus, shorter processes with varying thickness and less branching than type B. Type D (Ameboid); Big cell body merging with the processes, the nucleus fills out most of the cell body, and the processes are very short, thick and few. The average percentage of type A. B, C, and D of the total number of Iba1+ cells was calculated. Two-way ANOVA followed by Sidak’s multiple comparisons was applied.

### Single-cell suspension

Brain stems were harvested from animals sedated with ketamine and xylazine and thoroughly perfused with a minimum of 25 ml of PBS. Brain stems were cut into pieces and digested for 40 min at 37 °C with 5% CO_2_ in PBS supplemented with 1.6 IU ml^-1^ Collagenase/Dispase (Roche) and 0.5 mg ml^-1^ DNase I (Sigma-Aldrich). 20 μl of sterile EDTA was added to each sample and incubated at 37°C for another 5 min. Tissues were then mechanically disrupted through a 70um cell strainer into a single-cell suspension. Single-cell suspensions were washed with 25ml 2% FCS-HBSS (Thermo Fisher Scientific), centrifuged, and resuspended in a mixture of PBS with 0.5% BSA (VWR) and Debris Removal Solution (Miltenyi Biotec). After overlaying PBS with 0.5% BSA, suspensions were centrifuged, resuspended, and adjust the cell count to 10^6^ cells ml^-1^.

### Single-cell RNA-sequencing preparation

Single cell suspensions were converted to barcoded scRNAseq libraries following the manufacturer’s instructions of Chromium Single Cell 3’ Library, Gel bead & index kit and Chip G Kit (v3.1, 10x Genomics), aiming for recovering 10,000 cells per library. Quality control of libraries were performed with Tapestation 4200. After QC, libraries were sequenced on an MGI DNBSEQ-G400 sequencer.

### scRNA-seq data pre-processing

Reads demultiplexed from the DNBSEQ-G400 were mapped to a composite reference genome consisting of the mouse genome (GRCm39) and the HSV-1 genome (JX142173) using Cell Ranger v6.1 from 10x Genomics. Predicted doublets identified by the Python package solo (v1.0) were removed before transferring the unique molecular identifier (UMI) count matrix into a Seurat object using the R library Seurat (4.4.0) for downstream analysis. Cells with more than 200 detected genes and less than 30% mitochondrial gene content were considered qualified and retained. After quality control, the dataset comprised 143,915 cells and 29,390 featues. Normalization was performed using sctransform v2 on all samples merged by conditions. CCA integration was employed for batch effect correction to ensure consistency across batches. Dimensionality reduction using PCA and UMAP, followed by clustering with KNN, was sequentially performed on the integrated count matrix to simplify and visualize the dataset and to identify distinct cell clusters.

Differentially expressed genes (DEGs) across various conditions, cell types, and cell subpopulations were identified using the FindMarkers function after reverting SCT assays to corrected counts with the PrepSCTFindMarkers function.

### Viral transcripts analysis

The genomic copy number in cells containing viral transcripts was inferred from single-cell transcriptional profiles using the CopyKAT tool. CopyKAT utilizes depth information from the gene expression levels of adjacent genes to infer the genomic copy number in the corresponding region. The analysis was performed using the default parameters provided by CopyKAT.

### Cell type proportions analysis

To assess the dynamic changes in cell type proportions under various conditions, we employed statistical analyses to detect differences in cell type composition, which was conducted using the speckle package which uses generalized linear mixed models to model cell counts and performs permutation tests to assess the significance of observed differences in cell type proportions. The speckle package provides robust statistical tests to identify significant changes in cell type composition across different experimental conditions to accurately capture the dynamics of cell population shifts. All analyses were performed using the default settings of the speckle package.

### Functional gene modules analysis

DEGs with an adjusted p-value of less than or equal to 0.01 categorized as either upregulated or downregulated, were used for over-representation analysis against annotated gene sets. Gene Ontology analysis and KEGG Pathway analysis were conducted using R package clusterProfiler (v4.6.0). Additionally, Disease Ontology analysis was performed using DOSE on human gene ortholog sets derived from mouse DEG sets via the babelgene (v22.9) R package. All identified DEGs ranked by fold changes were inputted to gene set enrichment analysis.

Brain disease-related gene module analysis cited from BrainBase -CNCB-NGDC was conducted using SCPA (v1.5.4). SCPA assesses pathway activity by examining changes in the multivariate distribution of a given pathway across different conditions.

### Subclustering analysis

Subclustering was conducted using MultiK (v0.1.0) and SigClust to determine the optimal number of clusters, allowing for a more precise identification of distinct cell subpopulations. MultiK was utilized to evaluate the stability and consistency of the clusters across multiple resolutions (from 0.05 to 2.00 with a step size of 0.05). MultiK allowed us to assess cluster robustness and select the most stable clustering configuration. SigClust was used to statistically test the significance of the identified clusters, ensuring that they represented true biological variation rather than random noise. This comprehensive approach enhanced the resolution of our analysis, ensuring robust and reliable subclustering results and facilitating a deeper understanding of cellular heterogeneity within the populations.

### Condition distribution preference score

To evaluate the tissue specificity of each subpopulation, we employed a methodology based on the previous study Lineage Tracking Reveals Dynamic Relationships of T Cells in Colorectal Cancer. This approach involved analyzing the distribution of each subpopulation across different tissue types and comparing the observed cell counts to those expected by random distribution. Specifically, we computed the ratio of observed to expected cell numbers (Ro/e) for each subpopulation under various conditions. By calculating the Ro/e ratios, we identified subpopulations that were enriched or depleted in specific conditions. This analysis provided insights into the condition-specific roles and potential functional implications of each subpopulation in the context of viral infection.

### Comprehensive analysis of cellular development pathways in monocyte

We utilized an integrated analytical approach combining slingshot, RNA velocity analysis, Partition-based Graph Abstraction (PAGA) and SCENIC (v1.3.1) to unravel the cellular trajectories and regulated gene expression dynamics of monocyte.

Lineage reconstruction and pseudotime inference were performed using identified subclusters of monocytes to uncover the global structure using the dyno and slingshot packages. We applied RNA velocity analysis to capture the direction and pace of cellular transitions, introducing a temporal aspect to our trajectory insights and allowing us to forecast future cellular states from existing transcriptional profiles. Furthermore, we utilized PAGA to generate a graph-based abstraction, which provided a concise and informative visualization of the intricate cellular transitions and interactions. Additionally, SCENIC was used to infer gene regulatory networks and cell population specific transcriptional factors with high regulons activity scores, providing insights into the underlying regulatory mechanisms.

### Cell-cell communication network analysis

Differentially expressed and active ligand-receptor interactions between conditions were identified using the NicheNet and MultiNicheNet (v2.0.0) packages. NicheNet allowed us to pinpoint variations in cell-cell communication dynamics. To further elucidate the downstream effects, we inferred the signaling target genes of the communication pairs by assessing the enrichment of predicted target genes within the receiver cell types. This comprehensive approach enabled us to map out the intricate signaling pathways and understand the functional implications of these interactions in different cellular contexts.

## Acknowledgements

We thank Gitte Ulbjerg Toft for technical assistance in the laboratories and the animal facility at Department of Biomedicine for supporting mouse handling. Funding was received from The European Research Council (ERC-AdG ENVISION; 786602); The Novo Nordisk Foundation (NNF23OC0084931); The Lundbeck Foundation (R359-2020-2287); The Danish National Research Foundation (DNRF164). Danish Agency for Higher Education and Science through the Danish national research infrastructure CellX (5229-00009B). We would like to thank the Bioinformatics Core Facility (Jacob Høgfeldt), Department of Biomedicine, Aarhus University, for support and sparring. Thanks to GenomeDK and Aarhus University for providing computational resources and support that contributed to these research results.

## Author information

### Author contributions

The study was conceived and designed by L.S.R. and S.R.P. Animal experiments were designed by L.S.R. and S.R.P., and performed by X.L. and L.S.R. L.L. and L.S.R. optimized the protocol for preparation of brain single cell suspensions. S.R.N.J., M.M.N., and, K.T. were responsible for the GeoMx data acquisition and first data analysis. I.H.K. performed the in situ microglia morphology analysis and quantification of Iba1+ cells in tissue sections. X.D. was responsible for all the analysis of single cell and spatial transcriptomics data. Interpretation of data to develop the scientific story were performed by X.D., L.S.R., and S.R.P. with input from all authors. M.R-R., Y. L., L.L., L.S.R., and S.R.P supervised the work. Funding was acquired by S.R.P.. The figures were drafted by X.D. The manuscript was drafted by X.D., L.S.R., and S.R.P., and edited by all authors, who support the conclusions.

---

**Extended Data Fig. 1. Characterization of single-cell sequencing data from HSV-infected brain a**. Representative image of head swelling of mock or HSV-1 infected mice at day 4, 6 and 8 p.i. (n=10 pr. group). **b**, Graphical illustration of the localization of the areas in the brain stem and cerebellum subjected to GeoMx RNA analysis. **c,** ROI selection for GeoMx analysis of infected and control mice. **d**, Comparison of percentage of identified antisense transcripts in the single cell sequencing data from the individual mice. **e,** Proportion of variations in transcripts compared to reference genomic sequences. Each bar represents individual mice. **f,** Histogram with density curve of correlation between host gene expression and viral gene count at each time point post infection. Lower panel: Expression of Il1rn across cell types is shown as a violin plot. **g,** Violin plots of cell type markers of each annotated cell type. **h**, Differences in cell type proportions of the indicated annotated cell types over time in the dataset. **i,** Cell type count number per mouse of brain-resident ell types. Data were analysed using two-tailored one-way ANOVA.

**Extended Data Fig. 2. Pathway enrichment in different cell types in the HSV-1-infected mouse brain. a-f**, Enriched GO terms for up-regulated genes across different time points post-infection in the indicated cell type. **g,** Immunohistological staining with anti-CD3 on brain stem sections from mice infected for 5 and 8 days with HSV-1.

**Extended Data Fig. 3. Analysis of microglia subpopulations during HSV-1 infection. a,** Network plot illustrating the linkages between genes and biological concepts derived from enriched GO terms of up-regulated DEGs at various infection time points. **b,** Network plot illustrating the linkages between genes and biological concepts derived from enriched GO terms of down-regulated DEGs at various infection time points. **c,** Heatmaps of reported transcriptional marker genes for activated, ameboid, and ramified microglia in various microglia subpopulations. **d,** Comparison of enriched functional terms of up-regulated genes in subpopulations prevalent in control, day 6, and day 8 samples, respectively. **e,** Heatmap of reported marker genes for subpopulations of microglia, and associated phenotypes. **f,** Comparison of enriched function term of up-regulated genes in microglia subpopulations #1, #4, #15, and #16.

**Extended Data Fig. 4. Analysis of monocyte subpopulations in the HSV-1-infected mouse brain. a,** UMAP of dimensional reduction of monocyte subpopulations labeled by subclustering identity and sample time points respectively. **b,** Feature plots of marker genes of monocyte subtype. **c,** Dot plots showing activated and suppressed terms from GSEA analysis in subpopulation 4. The network plot (right) illustrating the linkages between genes and biological concepts derived from GSEA terms in subpopulation 4. **d,** Dot plots showing activated and suppressed terms from GSEA analysis in subpopulation 7. **e,** The network plot illustrating the linkages between genes and biological concepts derived from GSEA terms in subpopulation 10. **f,** UMAP labeled with RNA velocity of single cells, estimated by difference between unspliced and spliced mRNAs. **g,** PAGA map estimating directed connectivity using RNA velocity. **h,** Ridgeline plots of pseudotime index in each monocyte subpopulation.

**Extended Data Fig. 5. Analysis of endothelial cells in the HSV-1-infected mouse brain. a,** Plots of the most differentially expressed receptor-ligand communication pairs involving endothelial cells as receivers during the course of HSV-1 infection. **b,** Plots of the most differentially expressed receptor-ligand communication pairs between monocytes as sender cell and endothelial cells as receivers during the course of HSV-1 infection. **c,** UMAP of dimensional reduction of endothelial cell subpopulations labeled by subclustering identity and sample time points respectively. **d,** Heatmap of characteristic marker genes for endothelial cell subtypes and associated phenotypes. **e,** Dot plot of viral transcripts in subpopulations of endothelial cell. **f,** Dot plot showing the expression of pyroptosis, apoptosis and necrosis marker genes in endothelial cell subpopulations. g, overlay of Gsdme expression onto the UMAP of all annotated cell types in the single cell RNA sequencing dataset analysed in the study.

## Notes

### Competing Interest Statement

The authors have declared no competing interest.

