## Supplementary figures and images for "Temporally resolved single-cell RNA sequencing reveals protective and pathological responses during herpes simplex virus 1 CNS infection"

### Extended Data Fig 1

#### Extended Data Figure 1

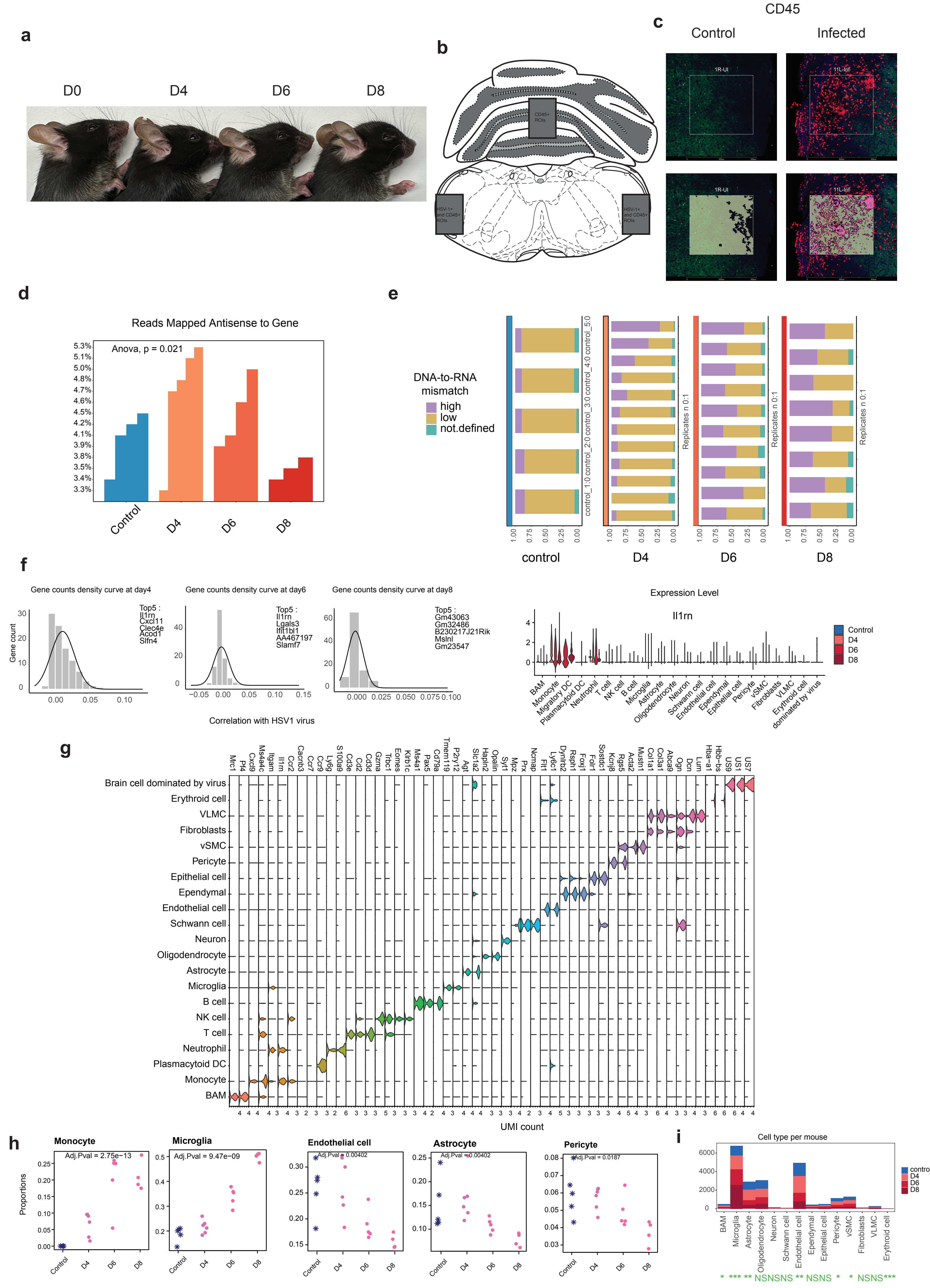

### Extended Data Fig 2

**Extended Data Figure 2**

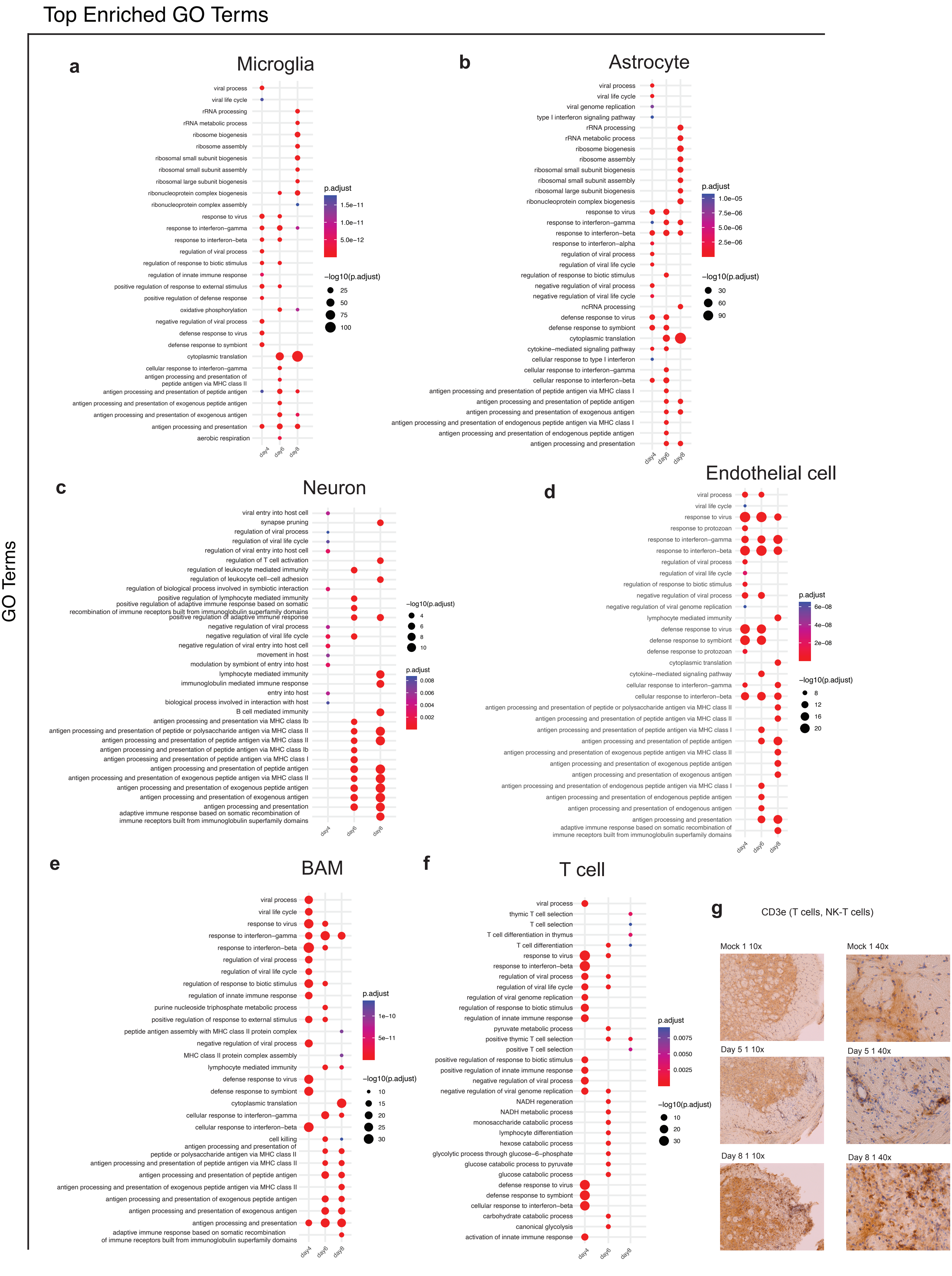

### Extended Data Fig 3

#### Extended Data Figure 3

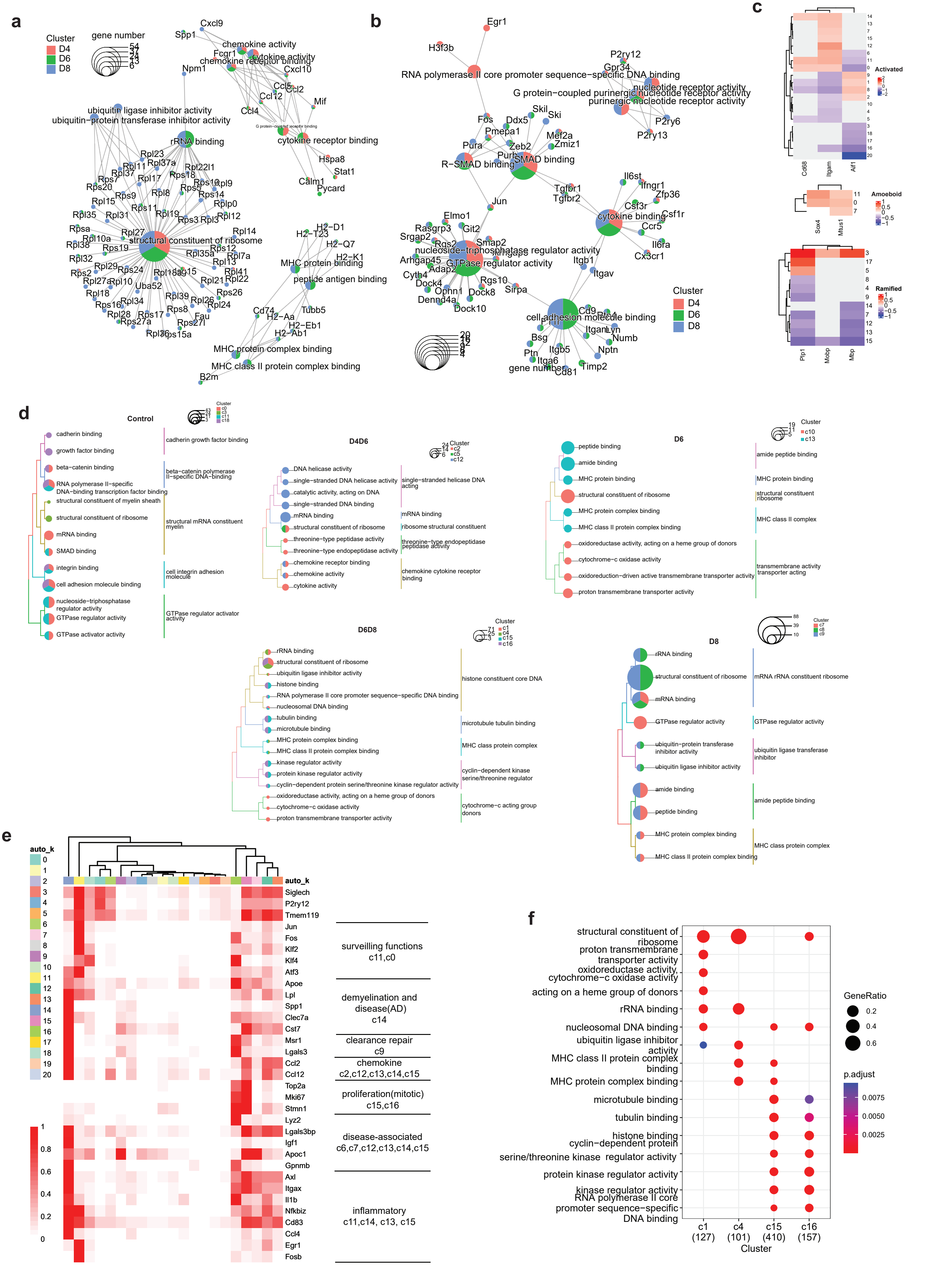

### Extended Data Fig 4

#### Extended Data Figure 4

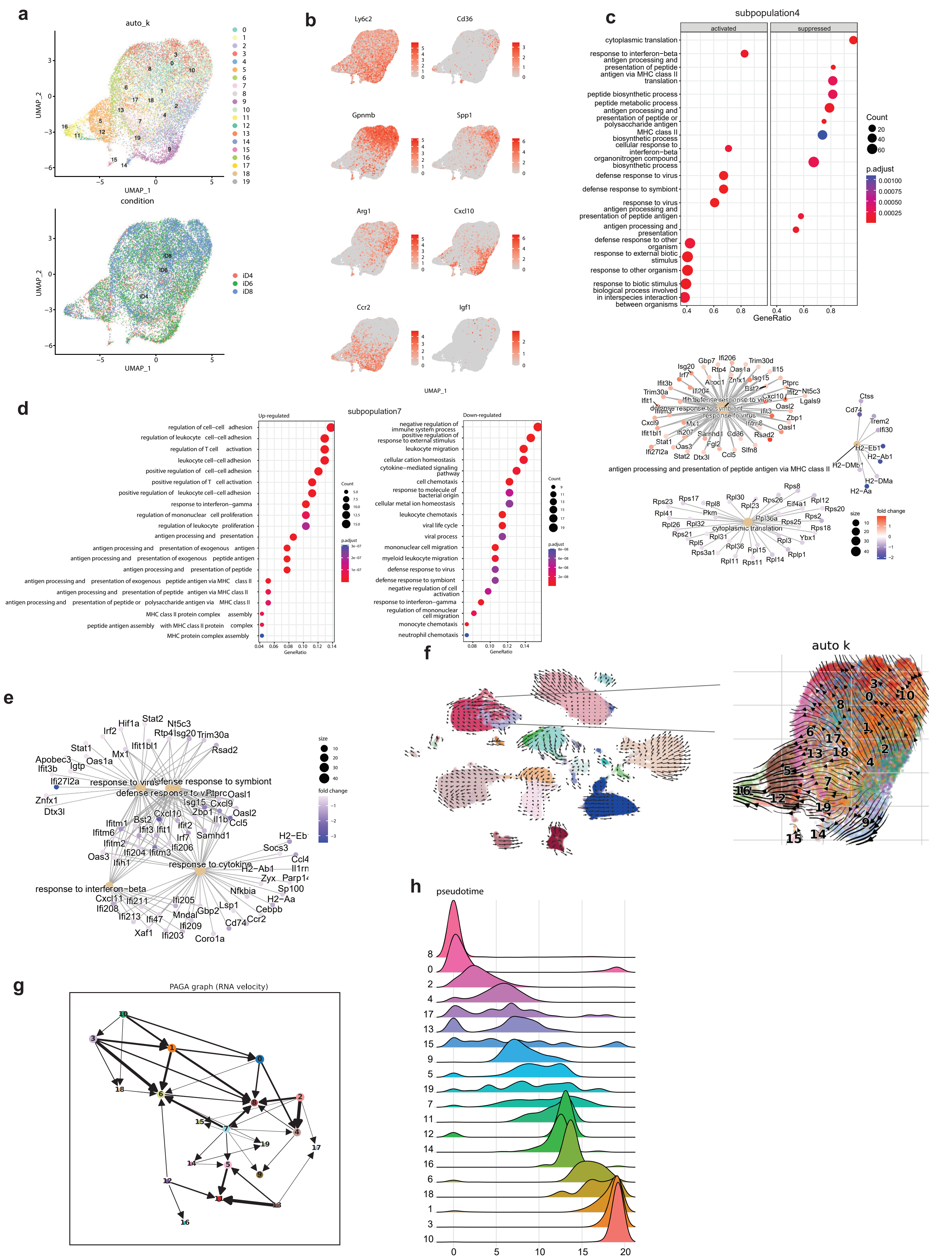

### Extended Data Fig 5

#### Extended Data Figure 5

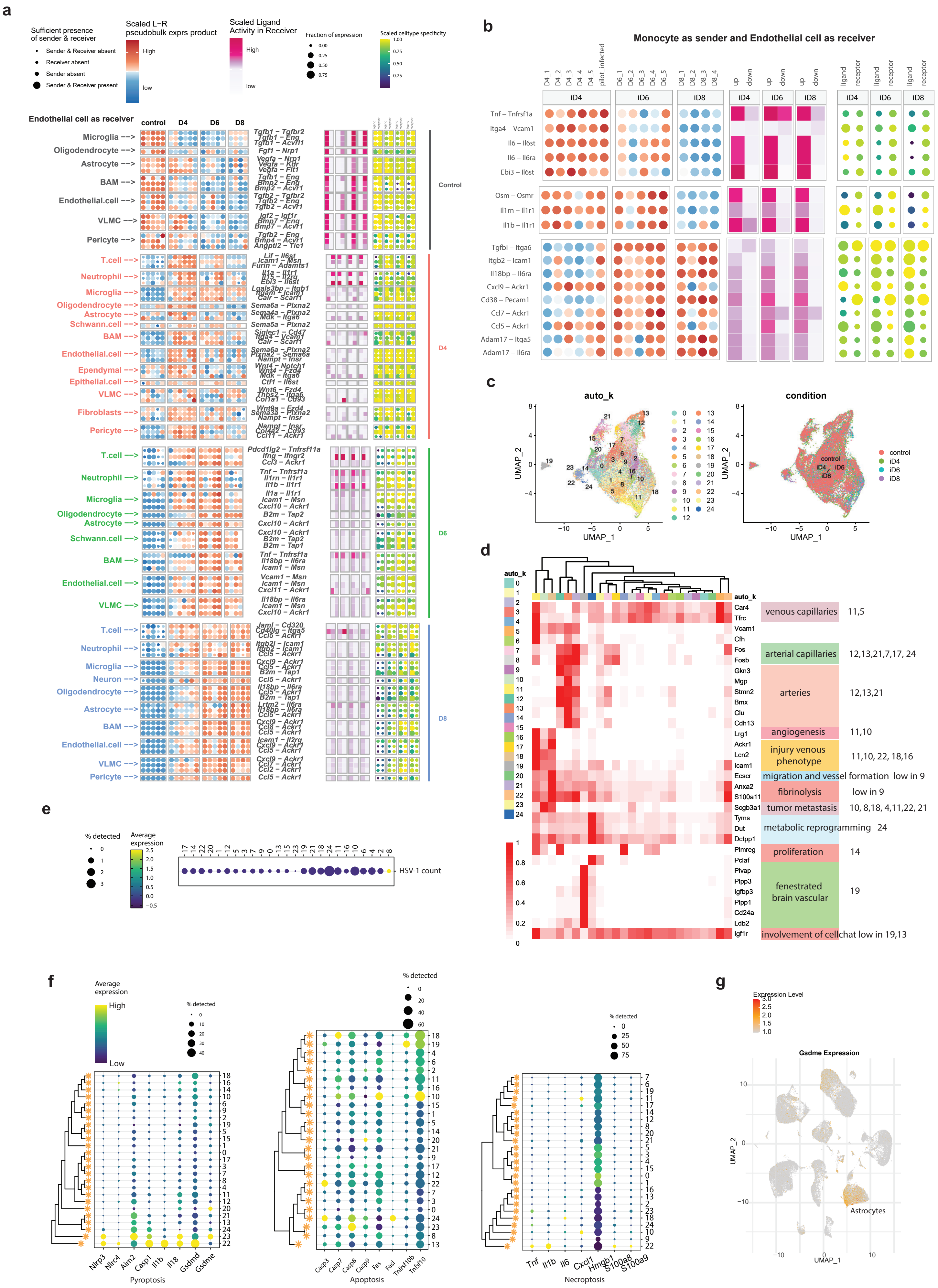
